# A novel stochastic simulation approach enables exploration of mechanisms for regulating polarity site movement

**DOI:** 10.1101/2020.11.30.404657

**Authors:** Samuel A. Ramirez, Michael Pablo, Sean Burk, Daniel J. Lew, Timothy C. Elston

## Abstract

Cells polarize their movement or growth toward external directional cues in many different contexts. For example, budding yeast cells grow toward potential mating partners in response to pheromone gradients. Directed growth is controlled by polarity factors that assemble into clusters at the cell membrane. The clusters assemble, disassemble, and move between different regions of the membrane before eventually forming a stable polarity site directed toward the pheromone source. Pathways that regulate clustering have been identified but the molecular mechanisms that regulate cluster mobility are not well understood. To gain insight into the contribution of chemical noise to cluster behavior we simulated clustering within the reaction-diffusion master equation (RDME) framework to account for molecular-level fluctuations. RDME simulations are a computationally efficient approximation, but their results can diverge from the underlying microscopic dynamics. We implemented novel concentration-dependent rate constants that improved the accuracy of RDME-based simulations of cluster behavior, allowing us to efficiently investigate how cluster dynamics might be regulated. Molecular noise was effective in relocating clusters when the clusters contained low numbers of limiting polarity factors, and when Cdc42, the central polarity regulator, exhibited short dwell times at the polarity site. Cluster stabilization occurred when abundances or binding rates were altered to either lengthen dwell times or increase the number of polarity molecules in the cluster. We validated key results using full 3D particle-based simulations. Understanding the mechanisms cells use to regulate the dynamics of polarity clusters should provide insights into how cells dynamically track external directional cues.

**Author summary:** Cells localize polarity molecules in a small region of the plasma membrane forming a polarity cluster that directs functions such as migration, reproduction, and growth. Guided by external signals, these clusters move across the membrane allowing cells to reorient growth or motion. The polarity molecules continuously and randomly shuttle between the cluster and the cell cytosol and, as a result, the number and distribution of molecules at the cluster constantly changes. Here we present an improved stochastic simulation algorithm to investigate how such molecular-scale fluctuations induce cluster movement across the cell membrane. Unexpectedly, cluster mobility does not correlate with variations in total molecule abundance within the cluster, but rather with changes in the spatial distribution of molecules that form the cluster. Cluster motion is faster when polarity molecules are scarce and when they shuttle rapidly between the cluster and the cytosol. Our results suggest that cells control cluster mobility by regulating the abundance of polarity molecules and biochemical reactions that affect the time molecules spend at the cluster. We provide insights into how cells harness random molecular behavior to perform functions important for survival, such as detecting the direction of external signals.

## Introduction

Cell migration, division, and differentiation require breaking the internal symmetry of the cell and establishing an axis of orientation. This symmetry breaking is referred to as polarity establishment. In eukaryotes, polarity establishment occurs as polarity factors, such as Rho-family GTPases, localize in a small region of the plasma membrane where they regulate the cytoskeleton to remodel cell morphology and generate motility [1]. In particular contexts, the polarity site can be highly dynamic. For example, migrating cells frequently change their direction of polarization as they navigate guided by changing environmental cues [2–4].

Polarity establishment has been well characterized in the budding yeast *Saccharomyces cerevisiae*. Yeast polarize in the contexts of budding and mating. The first step involves the clustering of the conserved master regulator of polarity, the Rho-GTPase Cdc42, at a site on the plasma membrane often referred to as the “polarity patch”. In the context of mating, detection of pheromone secreted by a potential mating partner can trigger polarization. However, the location of the initial polarity patch is inaccurate, and often misaligned with respect to the pheromone source [5,6]. The patch then relocates so that it is adjacent to a neighboring mating partner, allowing the two cells to fuse. Relocation of the polarity patch occurs in two stages. Initially the polarity patch is highly dynamic, rapidly assembling, disassembling, and moving along the cell membrane. In the next stage, Cdc42 organizes into a more concentrated patch with reduced mobility [5,6]. The initial rapid movement of the polarity patch is thought to be an exploratory phase to locate a mating partner. The remaining mobility of the patch during the second stage may be necessary to correct errors made during the exploratory phase. This view is supported by the observation that in experiments using externally imposed pheromone gradients, cells that did not polarize toward the gradient during the exploratory phase were able to reorient the polarity patch in the direction of the gradient [7–12]. Investigations combining experimental studies with mathematical modeling showed that actin-based vesicle delivery to the polarity patch is a key driver of patch movement during the second stage [7,13,14]. However, the mechanisms responsible for generating highly dynamic clustering during the exploratory phase and the transition to more stable polarity at the end of this stage are not well understood.

Recent studies revealed that the mobility of the polarity patch during mating is correlated with MAPK activity [5,6]. Pheromone-induced MAPK activity triggers polarization and drives changes in gene expression required for mating. During the early phases of mating when the polarity patch is highly mobile, MAPK activity is low. As the dynamic cluster of Cdc42 explores the membrane and relocates to a region near a mating partner, MAPK activity increases and the cluster of Cdc42 at the membrane becomes stable. Hegemann et al. [5] proposed that MAPK activity regulates patch mobility by inducing nuclear export of Cdc24, the GEF (activator) for Cdc42, thereby increasing Cdc42 activation at the membrane. They also proposed that stochastic fluctuations in the biochemical events underlying Cdc42 polarization drive the mobility of the cluster during the exploratory phase. The plausibility of the second claim was supported using a simple stochastic model for cell polarization adapted from [15]. Their mathematical formulation, however, did not address the mechanism for cluster stabilization.

To gain further insight into how chemical noise can induce cluster mobility and how cells can regulate cluster dynamics, we considered mechanistically detailed stochastic models of cell polarization. In a previous study, we used particle-based simulations to demonstrate that molecular-level fluctuations favor polarity establishment [16]. The stochastic simulations resulted in an extended parameter range over which polarity occurs and shorter times (1-5 min) for the emergence of a single polarity site in comparison to a deterministic reaction-diffusion version of the model. However, because particle-based simulations are computationally expensive, we were not able to address the stochastic behavior of the polarity patch over the time scales (10-100 min) associated with patch movement during yeast mating.

Particle-based simulations are computationally intensive because they track the position of every molecule in continuous space, and because simulating second-order reactions requires determining all pairs of molecules that are close enough to react. As an alternative, we represent the polarity circuit using the reaction-diffusion master equation (RDME), with individual simulations performed with an efficient spatial version of the Gillespie algorithm [17]. In this framework, space is discretized into grid units of finite size and molecules make diffusional jumps between adjacent grid units. Instead of tracking the exact position of every molecule, the RDME tracks the number of each chemical species within a grid unit, and reactions occur within grid units with a propensity proportional to the number of molecules and a rate constant usually referred to as “mesoscopic rate”. This approach can result in significantly reduced computation times; however, it does not represent accurately microscopic reaction-diffusion systems when association reactions are controlled by diffusion as in most polarity establishment models. Scale-dependent mesoscopic rates can more accurately reproduce microscopic dynamics in some diffusion-controlled systems [18–20]. However, we find that such mesoscopic rates are still inaccurate at the high molecular densities typical of the polarity system.

To overcome this limitation, we derived concentration-dependent mesoscopic rates inspired by the work of Yogurtcu et al [21]. They derived reaction rate constants that depend on the concentration to approximate diffusion-limited microscopic kinetics in homogeneous systems. We extended this method by making the mesoscopic rate constant dependent on the local concentration of the reactants within the grid units. We validated our approach in a 2D geometry by comparisons with particle-based simulations and applied it to study Cdc42 cluster dynamics accounting for molecular fluctuations. Key results were confirmed using full 3D particle-based simulations.

We found that molecular-level fluctuations can induce high mobility in Cdc42 clusters when clusters contain low numbers of the GEF for Cdc42 (the limiting polarity factor) and Cdc42 rapidly cycles between the cluster and the cytosol. Cluster stabilization was observed when GEF abundance increased or when the rate constant for association reactions between membrane-bound molecules increased. Accelerating such reactions stabilized clusters mainly by increasing the dwell time of Cdc42 or GEF molecules at the polarity patch. Interestingly, increasing the rate constant for formation of the Cdc42-GEF complex produced a switch-like transition in patch dynamics, suggesting its regulation may underlie patch stabilization during yeast mating.

## Results

### A. An improved approach for stochastic reaction-diffusion simulations

#### A.1. Background

We previously studied the effect of molecular-level fluctuations on the dynamics of polarity establishment using particle-based simulations [16]. In this approach, the Brownian motion of individual molecules occurs in continuous space and discrete time intervals of length Δ*t* (Fig 1A). Within any time interval Δ*t*, second-order reactions of the form A+B → C occur with probability λ Δ*t* if A and B are within a capture distance ρ, which is typically determined by the size of the reacting molecules. While particle-based simulations constitute a high-resolution representation of biochemical systems, they are computationally expensive, making it unfeasible to study systems with large number of molecules over long time scales.

**Figure 1.**
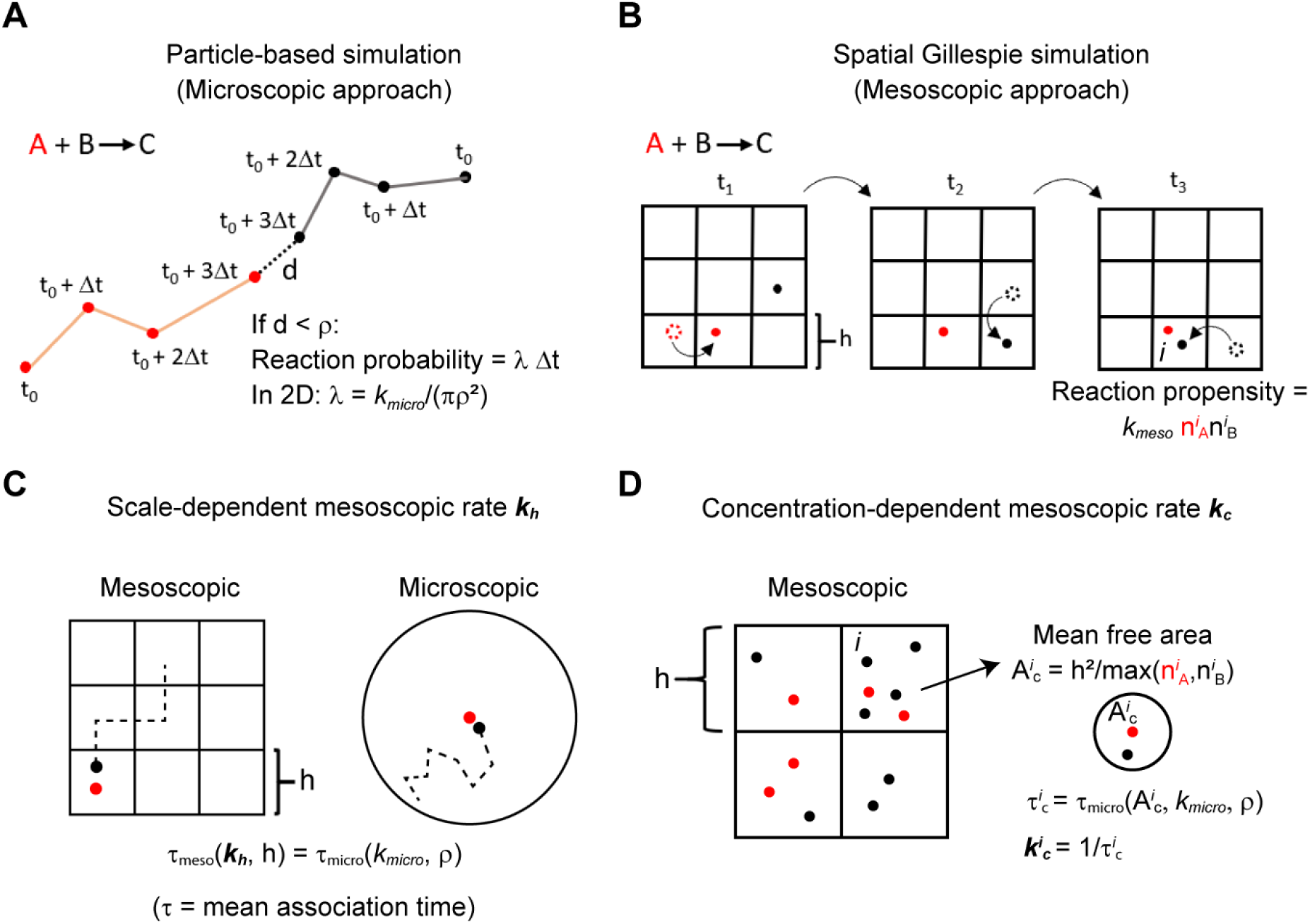
Schematic of the simulation of a generic bimolecular reaction A + B → C using a particle-based method and the spatial Gillespie approach. In the particle-based simulations (**A**) molecules undergo a random walk in continuous space and discrete time intervals Δ*t*. If a pair of reacting molecules are within a distance ρ characteristic of their size, they react with probability *λ* Δ*t* during an interval Δ*t*. In the spatial Gillespie approach (**B**), the domain is discretized using a grid, here with square elements of size *h*. Molecule jumps to adjacent grid elements and reactions take place sequentially at random times as a realization of the reaction-diffusion master equation. The propensity of a reaction within a grid element *i* is proportional to the number of molecules (n^*i*^_A_, n^*i*^_B_) and a mesoscopic rate constant *k*_meso_. (**C**) By setting *k*_meso_ equal to the scale-dependent mesoscopic rate *k_h_*, the mean association time of two molecules matches between the reaction-diffusion master equation and an analogous microscopic representation. (**D**) A concentration-dependent mesoscopic rate *k*^*i*^_c_ for the grid element *i* is estimated as 1/τ^*i*^_c_ where τ^*i*^_c_ is the mean association time in a microscopic representation of two molecules diffusing in a domain with area *A^i^_c_*. *A^i^_c_* is the mean free area between the most abundant reactant in the grid element *i* (further details in the main text and Methods section).

A computationally efficient alternative to particle-based modeling are simulations of the reaction-diffusion master equation (RDME) performed with an efficient spatial version of the Gillespie algorithm [17]. In RDME-based simulations, space is discretized into individual compartments, or grid elements, that can be occupied by multiple molecules (Fig 1B). Molecules can jump to adjacent grid elements with a propensity proportional to the diffusion coefficient (see Methods) and reactions occur with a propensity proportional to the molecular abundances of the reactants within the grid element and a mesoscopic rate constant *k*_meso_. Spatial Gillespie simulations are referred to as a mesoscopic approach in the sense that the size of the grid elements is normally larger than the size of a molecule (microscopic scale), but still significantly smaller than the whole system (macroscopic scale).

A problem with RDME simulations is that results depend on the spatial discretization and can deviate from more accurate microscopic models (e.g. particle-based simulations) especially when second-order reactions are controlled by diffusion [22]. This issue was addressed recently through the derivation of scale-dependent mesoscopic rate constants that ensure compliance with the microscopic kinetics in the limit of low molecular abundances [18–20]. Specifically, a generic system is considered consisting of a spatial domain containing two diffusing molecules that can undergo an association reaction. A mesoscopic rate is then derived using the requirement that the mean association time for the mesoscopic description is equivalent to that of the microscopic representation (Fig 1C, Methods). This “scale-dependent” mesoscopic rate constant depends on the grid element size *h*. If the association reaction is reversible, a mesoscopic dissociation rate is computed by ensuring that the equilibrium behavior of the mesoscopic description is identical to that of the microscopic representation.

In Section A.2. we evaluate the scale-dependent mesoscopic rate *k_h_* derived in [19,20] by looking at simulations of simple reaction schemes and comparing the results with particle-based simulations. We note that although *k_h_* yields accurate results at low concentrations, it shows significant deviations from particle-based simulations in scenarios with high molecular abundances. In Section A.3. we propose a concentration-dependent mesoscopic rate *k_c_* that accurately simulates systems over a broad range of concentrations. In Section A.4 we evaluate both *k_h_* and *k_c_* in a biochemical model for polarity establishment in yeast.

#### A.2. Evaluation of the scale-dependent mesoscopic rate *k_h_*

To benchmark spatial Gillespie simulations using *k_h_*, we compared their predicted kinetics with particle-based simulations. We initially focused on two prototypical reactions: irreversible association A+B → C and reversible association A+B ↔ C in diffusion-controlled regimes. As our goal is to model reaction and diffusion at the cell membrane, simulations are performed in a 2D domain with periodic boundary conditions.

We present timeseries of the mean number of molecules of one of the reactants in Fig 2, A-D. At low concentrations of reactants, simulations using *k*_h_ closely approximated results from particle-based simulations for both irreversible and reversible second-order reactions (Figs 2 A,B). At higher concentrations, however, the early kinetics of both the reversible and irreversible reactions showed significant deviations from the particle-based results (Figs 2 C,D). At later times the deviations are reduced as the number of molecules decreases. We note that it is not possible to achieve higher accuracies by reducing the size of the grid elements, because for this parameter regime, it is not possible calculate *k_h_* for *h* smaller than ~5ρ (Methods) [19,20].

**Figure 2.**
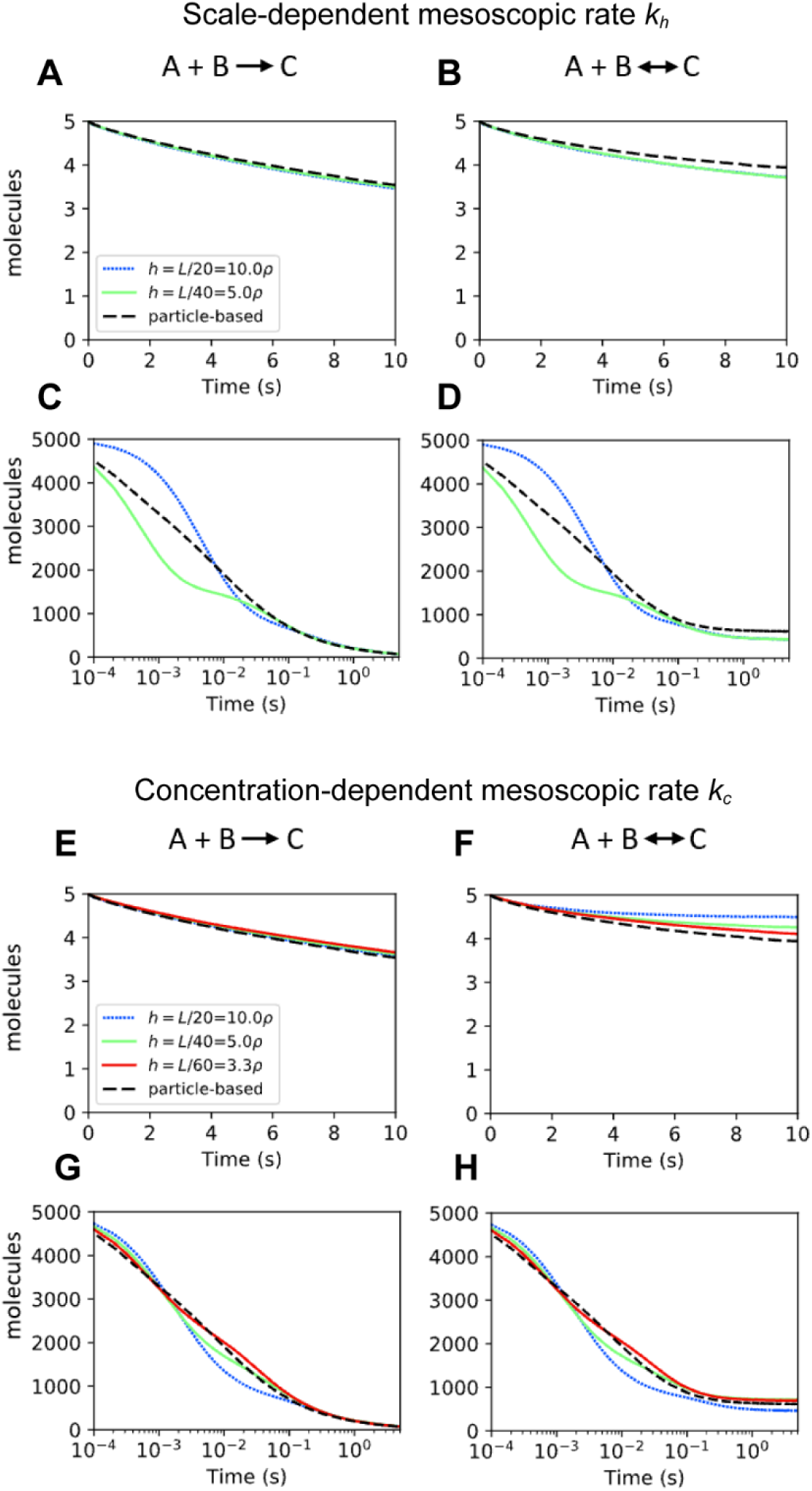
Spatial Gillespie simulations of the reactions A+B → C and A+B ↔ C using the mesoscopic rates *k_h_* and *k_c_* are benchmarked against particle-based results. We present the mean total number of species A as a function of time. **(A-D)** show spatial Gillespie simulations using the mesoscopic rate *k_h_* with initial low abundance of reactants in **(A, B)** (total A = total B = 5, total C = 0 at t=0) and initial high abundance (total A = total B = 5000, total C = 0 at t=0) in **(C-D)**. **(E-H)** show corresponding simulations to **(A-D)** but using the mesoscopic rate *k_c_*. In all the simulations, the degree of diffusion control is *k_micro_*/*D*_tot_ = 50, with *D*_tot_ = 2*D* and *D* = 0.0025μm^2^/s. The size of the domain is *L* = 1μm, ρ = 0.005μm. For the reversible reaction A+B ↔ C, the microscopic dissociation rate constant *k^d^_micro_* is 1/s in **(B, F)**, and 10/s in **(D, H)**.

We also computed timeseries of the standard deviation to quantify the fluctuations around the mean for each case in Figs 2 A-D (S1 A-D Figs). This metric also showed agreement between spatial Gillespie simulations using *k_h_* and particle-based simulations at low concentrations, and increased deviations at high concentrations at early times. Overall, the mesoscopic approach using *k_h_* provides a good approximation to the microscale dynamics for irreversible and reversible diffusion-controlled reactions only if the density of reactants is low.

To understand why *k_h_* showed accurate results for systems with low abundances but loses accuracy at high concentrations, remember that *k_h_* was derived to precisely reproduce the association time in a two-molecule system (Fig 1C, Methods). In a diluted system containing n pairs of reacting molecules, the expected time for an association event is similar to that in a set of n independent 2-molecule systems, and simulations using *k_h_* provide a good approximation of the kinetics. At high concentrations, however, molecules are more likely to associate with partners in their vicinity before diffusing over a significant portion of the spatial domain. Because the derivation of *k_h_* does not consider such multi-molecule effects, simulations lose accuracy.

#### A.3. A concentration-dependent mesoscopic rate *k*_c_ improves accuracy at high concentrations

To improve accuracy at high concentrations, we applied a concentration-dependent rate constant *k*_c_ = 1/τ_c_, where τ_c_ is the mean association time of two molecules within an effective area *A*_c_. *A*_c_ is the mean free area between molecules of the more abundant reactant in the grid element (Fig 1D, Methods) and is estimated as the grid element area *h*^2^ divided by the number of molecules of the more abundant reactant. This approach was inspired by the work of Yogurtcu et al [21], in which the authors estimated a concentration-dependent rate constant for spatially homogeneous systems. In our simulation approach, *k_c_* depends on the local concentration through *A*_c_ in each grid element, and therefore it changes in space and time as the system evolves.

For the case of reversible second-order reactions, a concentration-dependent mesoscopic dissociation rate *k*_c_^d^ is estimated in a similar way as for *k_h_*^d^. That is, the equilibrium behavior of a two-species system in the mesoscopic representation is matched to that of the microscopic formulation for a pair of molecules diffusing in a domain with area *A*_c_ (Methods).

Even though *k*_c_ was derived to provide accurate results at high concentrations, it also showed accurate results for low molecular abundances in the reaction A + B → C (Fig 2E). In the reversible reaction A+B ↔ C, simulations showed deviations that decreased with smaller grid element size *h* (Fig 2F). At high molecular abundance, *k*_c_ showed increased accuracy compared to *k_h_* in both irreversible and reversible reactions (compare Figs 2 C,D and 2 G,H). The increase in accuracy using *k_c_* was also observed in the fluctuations around average concentrations (S1 C,D and S1 G,H Figs).

#### A.4 Evaluation of mesoscale simulations in a model of the yeast polarity circuit

We next evaluated the mesoscale rates *k_h_* and *k_c_* in a reaction-diffusion model for polarity establishment in budding yeast adapted from [23] (Fig 3A). Central to this biochemical network is the Rho-GTPase Cdc42. Cdc42 can exist in an inactive (GDP bound) form Cdc42D that shuttles between the membrane and the cytosol, and an active (GTP bound) form Cdc42T that localizes to the cell membrane. Deactivation occurs as Cdc42 hydrolyzes GTP into GDP, a process accelerated by GTPase activating proteins (GAPs). GAP activity is considered implicitly using a pseudo-first order deactivation rate constant. The activation of Cdc42 is catalyzed by the guanine nucleotide exchange factor Cdc24 (GEF) that binds Cdc42D and facilitates the exchange of GDP for GTP. GEF molecules can shuttle between membrane and cytosol, and at the membrane they activate Cdc42. Once GEF binds Cdc42D, it activates the Rho-GTPase and dissociates from the resulting Cdc42T in a single step. There is positive feedback in the levels of active Cdc42 because Cdc42T can bind a GEF molecule forming a complex Cdc42T-GEF that can activate neighboring Cdc42D molecules. In this model Cdc42T can recruit both membrane-bound and cytosolic GEF molecules. In the following simulations, membrane and cytosol are represented as coincident 2D square domains. What distinguishes the membrane and cytosol are the diffusion coefficients for the molecular species in each domain. The parameters were adapted from a 3D macroscopic model [13,23] into a 2D microscopic representation (Methods) and are presented in Table 1.

**Figure 3.**
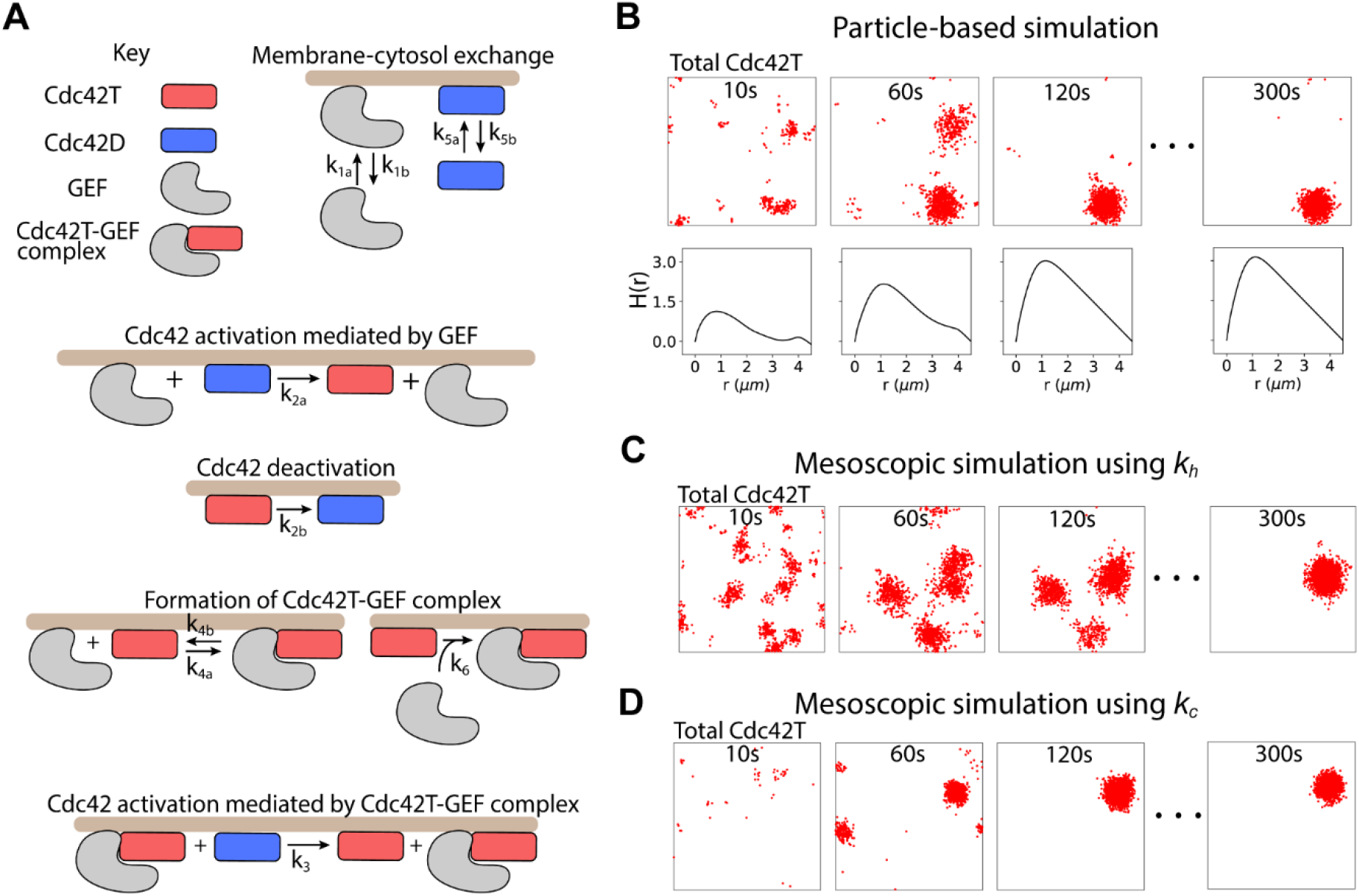
Cdc42 polarization in yeast simulated with particle-based simulations and spatial Gillespie simulations using the mesoscopic rates *k_h_* and *k_c_*. **(A)** Reactions in a model for polarization in budding yeast. Species that are not bound to the membrane (brown), dwell in the cytosol. Membrane and cytosol are represented in the simulations as juxtaposed 2D squared domains. **(B)** Time series of a particle-based simulation of the polarization model in **(A)**. In each snapshot, red dots show the positions on the membrane of all active Cdc42 molecules (Cdc42T and Cdc42T-GEF). The lower panels in **(B)** show a quantification of active Cdc42 clustering using the H(*r*) function (see Methods for details). **(C, D)** Spatial Gillespie simulations using *k_h_* **(C)** and *k_c_* **(D)** with the same model parameters as the particle-based simulation in **(A)** and grid element size h = 5ρ. Pseudo-coordinates for each molecule are randomly sampled from the containing grid element and displayed as a red dot to facilitate comparison with particle-based simulations. Model parameters are presented in Table 1.

**Table 1.** Model parameters

| Description | Param. | 2D Model | 3D Model |
| --- | --- | --- | --- |
| $\text{GEF}_c \rightarrow \text{GEF}_m$ | $k_{1a}$ | $0.1 \text{ s}^{-1}$ | $0.07522 \mu\text{m s}^{-1}$ |
| $\text{GEF}_m \rightarrow \text{GEF}_c$ | $k_{1b}$ | $10 \text{ s}^{-1}$ | $10 \text{ s}^{-1}$ |
| $\text{Cdc42D}_m + \text{GEF}_m \rightarrow \text{Cdc42T}$ | $k_{2a}$ | $0.032 \mu\text{m}^2\text{s}^{-1}$ | $0.032 \mu\text{m}^2\text{s}^{-1}$ |
| $\text{Cdc42T} \rightarrow \text{Cdc42D}_m$ | $k_{2b}$ | $0.63 \text{ s}^{-1}$ | $0.63 \text{ s}^{-1}$ |
| $\text{Cdc42D}_m + \text{Cdc42T-GEF} \rightarrow \text{Cdc42T}$ | $k_3$ | $0.07 \mu\text{m}^2\text{s}^{-1}$ | $0.07 \mu\text{m}^2\text{s}^{-1}$ |
| $\text{GEF}_m + \text{Cdc42T} \rightarrow \text{Cdc42T-GEF}$ | $k_{4a}$ | $0\text{-}2 \mu\text{m}^2\text{s}^{-1}$ | $0\text{-}2 \mu\text{m}^2\text{s}^{-1}$ |
| $\text{Cdc42T-GEF} \rightarrow \text{GEF}_m + \text{Cdc42T}$ | $k_{4b}$ | $10 \text{ s}^{-1}$ | $10 \text{ s}^{-1}$ |
| $\text{Cdc42D}_c \rightarrow \text{Cdc42D}_m$ | $k_{5a}$ | $4 \text{ s}^{-1}$ | $3.009 \mu\text{m s}^{-1}$ |
| $\text{Cdc42D}_m \rightarrow \text{Cdc42D}_c$ | $k_{5b}$ | $6.5 \text{ s}^{-1}$ | $6.5 \text{ s}^{-1}$ |
| $\text{GEF}_c + \text{Cdc42T} \rightarrow \text{Cdc42T-GEF}$ | $k_6$ | $0.2 \mu\text{m}^2\text{s}^{-1}$ | $0.15 \mu\text{m}^3\text{s}^{-1}$ |
| $\text{Cdc42D}_c + \text{Cdc42T-GEF} \rightarrow \text{Cdc42T}$ | $k_7$ | $0.5 \mu\text{m}^2\text{s}^{-1}$ | $0.376 \mu\text{m}^3\text{s}^{-1}$ |
| Diffusion coefficient in cytoplasm | $D_{\text{cyto}}$ | $10 \mu\text{m}^2\text{s}^{-1}$ | $10 \mu\text{m}^2\text{s}^{-1}$ |
| Diffusion coefficient on membrane | $D_{\text{memb}}$ | $0.0045 \mu\text{m}^2\text{s}^{-1}$ | $0.0045 \mu\text{m}^2\text{s}^{-1}$ |
| Membrane surface area | $A_m$ | $64 \mu\text{m}^2$ | $64 \mu\text{m}^2$ |
| Total Cdc42 | Cdc42 | 5000 molecules | 5000 molecules |
| Total GEF | GEF | 15-700 molecules | 15-700 molecules |
| Membrane thickness | $Th$ | $0.0083 \mu\text{m}$ | $0.0083 \mu\text{m}$ |
| Cell volume | $V_c$ | $48.144 \mu\text{m}^3$ | $48.144 \mu\text{m}^3$ |
| Membrane volume | $V_m$ | $0.53 \mu\text{m}^3$ | $0.53 \mu\text{m}^3$ |
| $V_m/V_c$ | $\eta$ | 0.011 | 0.011 |
| Reactive radius | $\rho$ | $0.02 \mu\text{m}$ | $0.02 \mu\text{m}$ |

Particle-based simulations initialized with all Cdc42 and GEF molecules randomly located in the cytosol evolved into a polarized distribution of total Cdc42T (Cdc42T and Cdc42T-GEF) at the membrane (Fig 3B) [16]. At early times, small fluctuations in the levels of Cdc42T are amplified by positive feedback reactions resulting in growing clusters of the active Rho-GTPase (Fig 3B 10 s). Several clusters can form that compete for polarity factors in the cytosol until just one cluster remains (Fig 3B, 60 s - 120 s). The remaining cluster grows, depleting polarity factors in the cytosol, until a steady state is reached when the inward and outward fluxes of molecules to the cluster balance (Fig 3B 120 s – 300 s).

In Figs 3C and 3D we present snapshots of representative mesoscopic simulations using the scale-dependent (*k_h_*) and concentration-dependent (*k_c_*) rates, respectively. To facilitate comparison with particle-based simulations, spatial pseudo-coordinates for each molecule were obtained by randomly sampling within the grid element containing the molecule. Mesoscopic simulations using both *k_h_* and *k_c_* reached a steady-state polarity cluster (Figs 3 C,D 300 s) with a size comparable to that of the particle-based simulation in Fig 3A. There were differences, however, in the time evolution of polarization between the different methods. The simulation using *k_h_* took longer to evolve into a single polarity cluster, while the simulation using *k_c_* polarized faster compared to the particle-based approach. We note that these simulations were run using a grid element *h* = 5ρ which corresponds to the finest grid possible using *k_h_*.

To quantify the dynamics of polarization we measured clustering of total Cdc42T with the function H(*r*) (Fig 3B lower panels) [16,24,25] (Methods). H(*r*) has the desired properties that H(*r*) = 0 for a random distribution of molecules, and for a clustered distribution, H(*r*) shows a maximum at a value *r* = *r_max_* which provides a measure of the cluster size. H(*r*) showed a maximum at *r* = 1.1 μm that increased over time (Fig 3B lower panels). We therefore chose H(*r* = 1.1 μm) as a metric for polarization. We note that quantifying clustering with max H(*r*) does not qualitatively change the results. In Fig 4A we show the time evolution of the mean of H(*r* = 1.1 μm) over different realizations for the different simulation methods. The shading indicates standard deviation to illustrate the variability in polarization dynamics. At early times, simulations using *k_h_* matched particle-based simulations but showed deviations at later times, taking longer to polarize. On the other hand, simulations using *k_c_* displayed overall faster polarization than the other methods. By defining the polarization time as the moment when the standard deviation stabilizes, it is apparent that simulations using *k_h_* have a longer time of polarization (240 s) compared to particle-based simulations (180 s), while using *k_c_* results in faster polarization (130 s).

**Figure 4.**
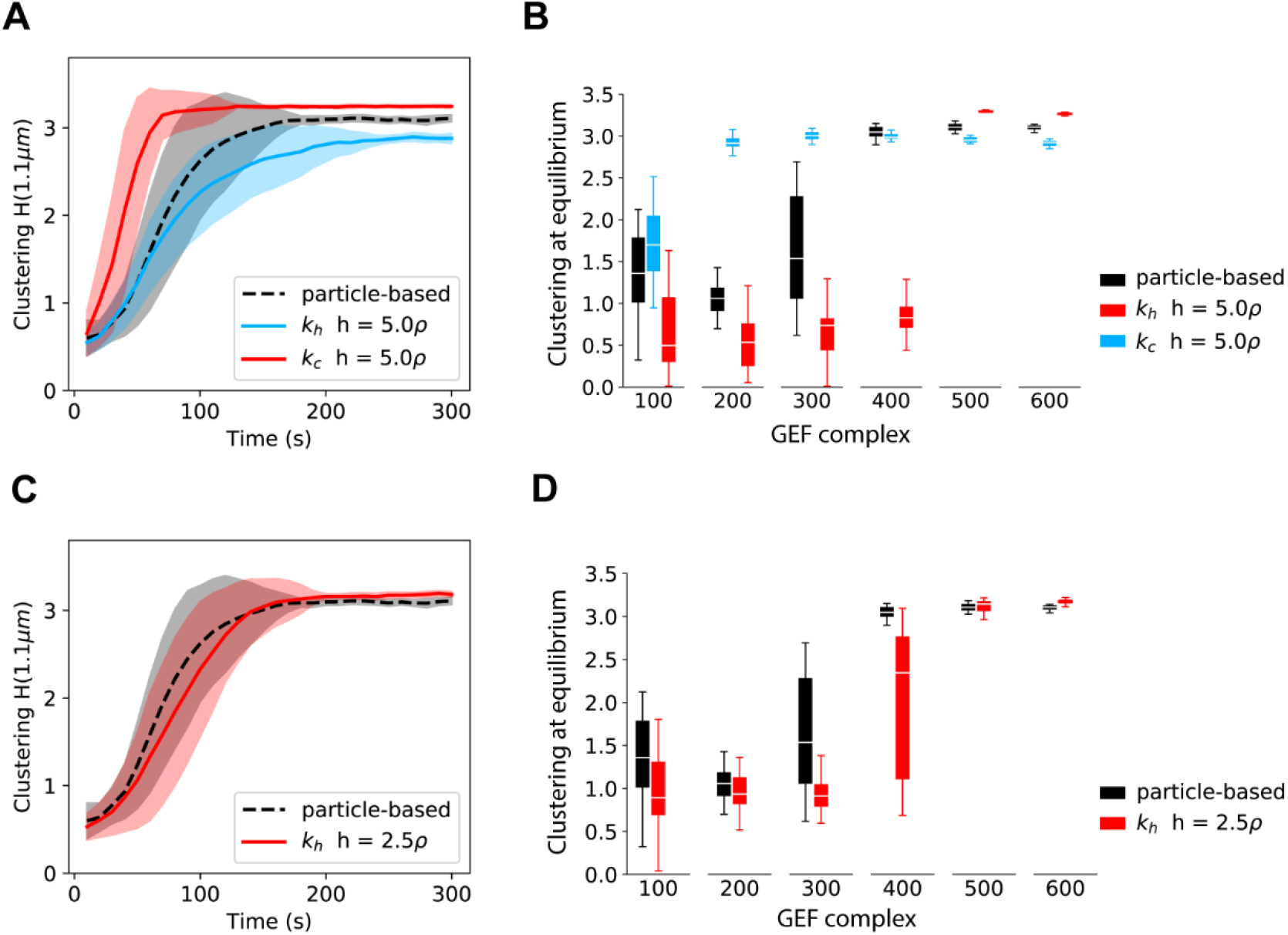
Spatial Gillespie simulations using the mesoscopic rates *k_h_* and *k_c_* are benchmarked against particle-based simulations by looking at Cdc42 polarization dynamics and equilibrium. **(A)** Time evolution of the clustering of total active Cdc42 from simulations of the polarity model in Fig 3A with parameters in Table 1. We contrast results from particle-based simulations and the spatial Gillespie approach using either *k_h_* or *k_c_* and a grid element size *h* = 5ρ. Clustering at a particular timepoint is quantified as the mean of H(*r* = 1.1 μm) over 30 simulations. Uncertainty intervals are computed as mean ± standard deviation and presented as a shaded region enveloping the mean. **(B)** Clustering at equilibrium from simulations in **(A)** for different values of the total amount of GEF. The clustering at equilibrium is computed as the mean of H(*r* = 1.1 μm) between 250s and 300s. The box plots were generated with the clustering at equilibrium from 30 simulations. **(C)** and **(D)** are corresponding figures to **(A)** and **(B)** respectively, the only difference is that the spatial Gillespie simulations are run with *h* = 2.5ρ and using only *k_c_* as *k_h_* cannot be computed for such *h* in this model (see Methods).

We further characterized the simulation approaches by looking at the equilibrium behavior for different amounts of available GEF molecules (Fig 4B). For high amounts of GEF all the methods showed polarization, but as the number of GEF molecules was reduced, simulations remained in a non-polarized state. While particle-based simulations efficiently polarized for GEF amounts greater than 300, mesoscopic simulations using *k_h_* polarized for numbers of GEF greater than 100, showing polarization in a regime where the microscopic simulations do not spontaneously polarize. On the other hand, mesoscopic simulations using *k_c_* showed polarization for GEF amounts greater than 400, failing to polarize at a value of 400 GEF where the microscopic approach shows polarization.

As the mesoscopic simulations with both *k_h_* and *k_c_* presented in Figs 4A and 4B showed discrepancies with respect to particle-based simulations, we sought to obtain more accurate results by reducing the grid element size, which so far was set to *h* = 5ρ. We were able to do this only for simulations with the concentration-dependent rate *k_c_*, because for the parameters of the model, *k_h_* cannot be computed for grid elements smaller than 5ρ (see Methods). For *h* = 2.5ρ we observed a significant improvement in accuracy using *k_c_* both in the dynamics of polarization (Fig 4C) and in the equilibrium behavior for different values of the number of GEF molecules (Fig 4D).

The increase in accuracy using *h* = 2.5ρ came with a cost of ≈ 5X increase in computation time with respect to simulations using *h* = 5ρ. Particle based simulations, on the other hand, were ≈ 10X more computationally expensive with respect to mesoscopic simulations using *h* = 5ρ. We note that particle-based simulations were performed with the highly optimized simulation platform Smoldyn [26,27], whereas mesoscopic simulations were run using our own custom written C code, which has not been optimized for computational performance.

### B. Mechanisms regulating mobility of the polarity site

#### B1. Requirements for high patch mobility

When yeast cells are presented with pheromone from a potential mating partner, Cdc42 forms dynamic clusters that explore the membrane for 10-100 min before stabilizing in a region close to the pheromone source [5,6]. To efficiently and accurately characterize polarity cluster dynamics over physiologically relevant timescales, we used spatial Gillespie simulations with concentration-dependent rates *k_c_* for bimolecular reactions.

To assess the mobility of the polarity cluster we tracked the centroid of the distribution for active Cdc42 over time (Fig 5A). To illustrate patch movement, we marked the initial position of the patch centroid using a green dot and its current position using a black dot. With our initial parameterization (Table 1), however, the patch did not move significantly over a 60 min time interval. It has been suggested that the amount of GEF in the cytosol strongly regulates cluster dynamics [5]. To investigate this effect, we ran simulations with different GEF levels, computed the mean squared displacement (MSD) of the patch centroid and used these values to estimate an effective diffusion coefficient (*D_patch_*) for the patch (Fig 5B). Decreasing GEF abundance increased patch mobility, but the system lost polarity at moderate GEF levels (≈ 450 molecules) before patch movement increased substantially. Therefore, we investigated if varying any of the rate constants would allow the system to polarize at lower GEF abundances. After testing all the reactions in the model, we observed that the following modifications allowed the system to polarize with low GEF abundances (100 or less molecules): 1) decreasing the rate constant *k_2b_* for Cdc42T deactivation, 2) decreasing the rate constant of dissociation of the Cdc42T-GEF complex *k_4b_*, 3) increasing the rate constant *k_5a_* for membrane binding of cytosolic Cdc42D and 4) increasing the rate constant *k_6_* for association of Cdc42T with cytosolic GEF to form the Cdc42T-GEF complex (S2 Fig). We computed patch mobility in simulations where both *k_2b_* and *k_5a_* where modified, which resulted in polarization down to 15 GEF molecules (Figs 5 C,D) and in simulations where *k_6_* was increased (Figs 5 E,F). Interestingly, as the total GEF abundance was decreased, only increasing *k_6_* generated highly mobile patches (compare Figs 5D and 5F which are representative realizations from the points indicated by the red arrows in Figs 5C and 5E).

**Figure 5.**
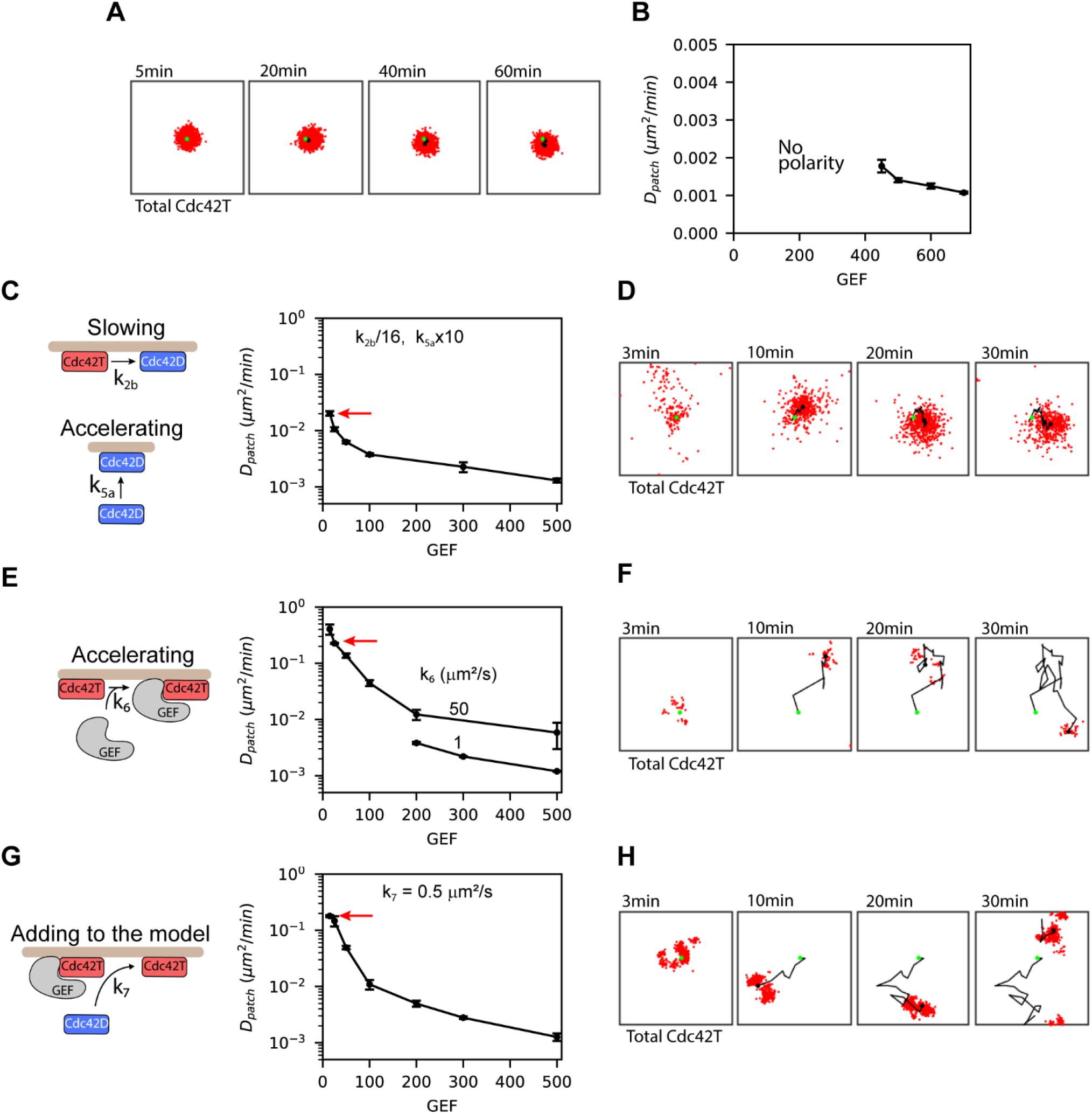
Direct recruitment of polarity factors to the patch enables highly mobile clusters at low GEF abundances. **(A)** Snapshots of the distribution of total active Cdc42 over a 1hr simulation. The black dot in each frame is the centroid of the polarity cluster, and the green dot is the centroid when the polarity cluster first formed. **(B)** Diffusivity of the centroid of the polarity patch (*D_patch_*) as a function of the total amount of GEF molecules in the simulations. *D_patch_* was obtained from the mean squared displacement (MSD) of the patch centroid by fitting the equation MSD(Δ*t_i_*) = 4*D_patch_* Δ*t_i_^β^* to the data, where Δ*t_i_* is a particular time interval, and β reflects the degree of anomalous diffusion. **(C)** Patch diffusivities as a function of total GEF for simulations where *k_2b_* has been decreased by a factor of 1/16 and *k_5a_* has been increased by a factor of 10 relative to the parameters in Table 1. For these simulations β ≈ 1. **(D)** Snapshots from a representative simulation in **(C)** as indicated by the red arrow. **(E)** Patch diffusivities as a function of total GEF for simulations where *k_6_* has been increased to either 1 μm^2^/s or 50 μm^2^/s. With *k_6_* = 1 μm^2^. Is polarization is lost when the number of GEFs is below 200. With *k_6_* = 50 μm^2^/s the simulations show polarization at even lower GEF amounts, in this case, β varied between 0.85 and 1. **(F)** Snapshots from a representative simulation in **(E)** as indicated by the red arrow. **(G)** Patch diffusivities as a function of total GEF after adding Reaction 7 to the model. β values were ≈ 0.85 for the two data points with highest mobilities, and close to 1 for the other points. **(H)** Snapshots from a representative simulation in **(G)** as indicated by the red arrow. Error bars for patch centroid diffusivities are standard errors from the least-squares fit use to compute *D_patch_*.

The rate constant *k_6_* governs the direct recruitment of cytosolic GEF to the patch through complex formation with Cdc42T. This reaction can occur rapidly, because diffusion in the cytosol is fast compared to diffusion in the membrane. Another potentially fast reaction not considered in the model is the recruitment of cytosolic Cdc42D directly to the patch. In cells, most inactive Cdc42 molecules are found in the cytosol bound to GDI proteins that hold them in their inactive state. In our model, for a cytosolic Cdc42D molecule to be activated it must first be inserted in the membrane, a step implicitly representing dissociation from the GDI protein. Once at the membrane Cdc42D has to laterally diffuse to react with a GEF. However, prior work has suggested that GEFs may displace Rho-GTPases from their GDI proteins [28,29]. Based on this observation, we updated our model to include a reaction where Cdc42T-GEF recruits and activates cytosolic Cdc42D (Fig 5G). An analogous reaction was included in a model for polarization by Klunder et al. (2013). Using a rate constant comparable to the one employed by Klunder et al. (2013) and maintaining the original values of the other rate constants, we observed high mobility of the polarity patch for low GEF abundances (Figs 5 G,H). Our results support the hypothesis that direct recruitment of fast diffusing cytosolic polarity factors to the patch promotes mobility of the polarity cluster.

To gain insight into the mechanism that generates a dynamic patch, we investigated several different patch properties. We first observed that the models shown in Figs 5C and 5G, which have substantially different patch mobility at low GEF abundance (replotted in Fig 6A on a log-log scale as “High mobility” and “Low mobility” models), have comparable amounts of total active Cdc42 and Cdc42T-GEF (the predominant state of GEF molecules at the patch) (Figs 6 B,C). The fluctuations in the total amount of active Cdc42 and Cdc42T-GEF, quantified as the coefficient of variation (*CV*) were overall comparable as well (Fig 6 D,E). We then evaluated the dwell time of Cdc42T at the membrane (see Methods) for the high and low mobility models (Fig 6F). The dwell time of Cdc42T was shorter in the high mobility model, indicating a faster cycling of Cdc42 between the cluster and the cytosol as compared to the low mobility model. We also observed that the shape of the patch fluctuates strongly in the high mobility model at low GEF levels (Fig 6G), while the low mobility model shows a more consistent distribution (Fig 6H). We quantified the variation over time in the distribution of Cdc42T (*CV_patch_*) (see Methods) and observed that the behavior of *CV_patch_* is qualitatively similar to that of *D_patch_* (Fig 6I). In fact, a plot of *D_patch_* and *CV_patch_* with data from the two models shows that these two quantities are highly correlated (Fig 6J). These results indicate that patch mobility is correlated with spatial fluctuations in the Cdc42T distribution, rather than average fluctuations in the total abundance of Cdc42T or its GEF at the polarity patch. The fluctuations in the Cdc42T distribution are stronger when GEF abundance is low and Cdc42 cycles rapidly between the cluster and the cytosol.

**Figure 6.**
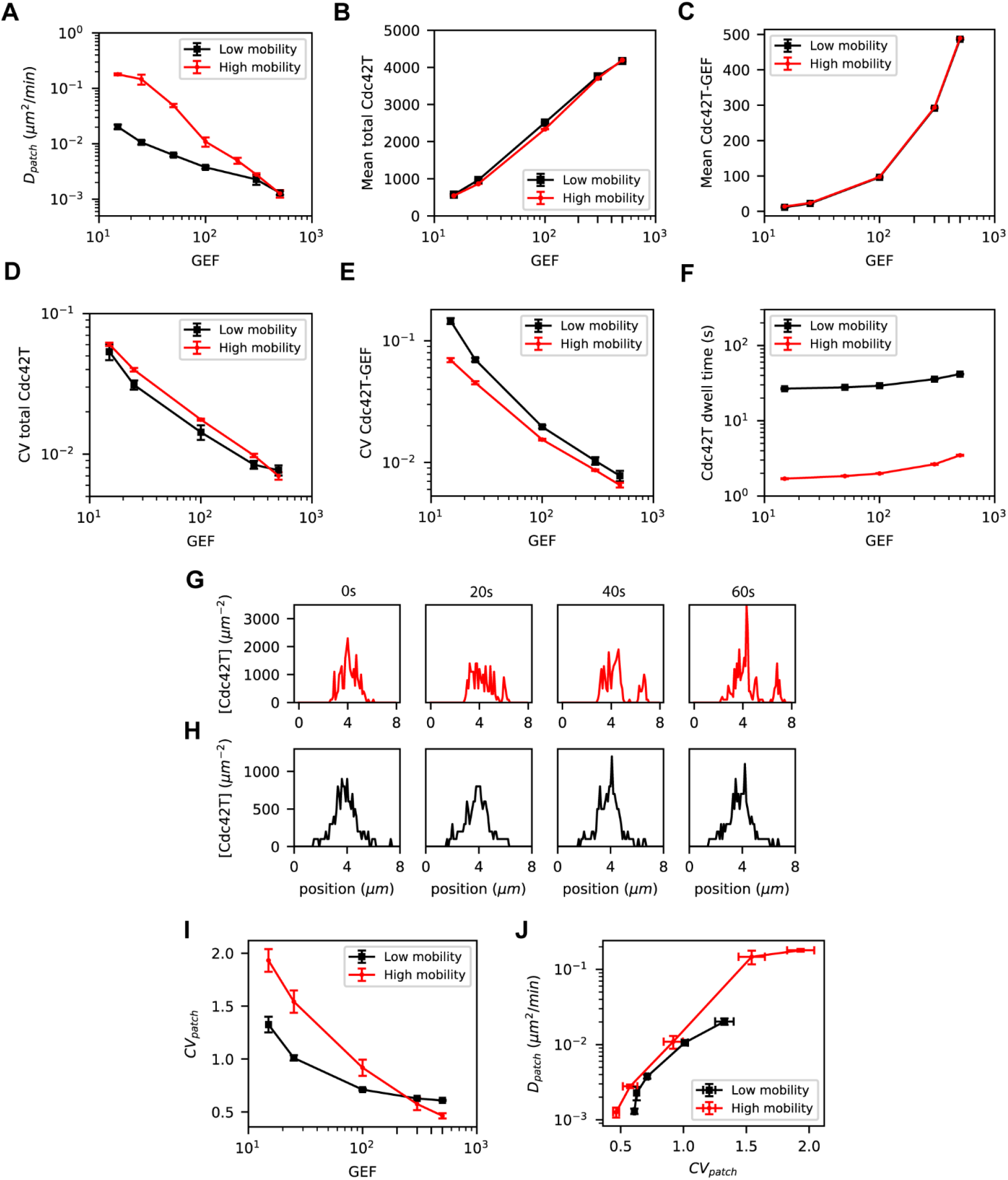
Cluster mobility correlates with rapid fluctuations in the distribution of active Cdc42 and is facilitated by short dwell times of Cdc42 at the patch and low GEF abundances. Comparison of different metrics as a function of total GEF molecules for a “Low mobility” model (from Fig 5C) where *k_2b_* has been decreased X/16 and *k_5a_* has been increased 10X, and a “High mobility” model (from Fig. 5G) that includes Reaction 7. **(A)** Patch centroid diffusivities, **(B)** Mean number of total Cdc42T, **(C)** coefficient of variation (*CV*) of total Cdc42T, **(D)** mean number of Cdc42T-GEF, **(E)** *CV* of Cdc42T-GEF, **(F)** dwell time of Cdc42T at the cluster, **(G)** Snapshots of the lateral profile of the concentration of total Cdc42T molecules for the High mobility model and **(H)** Low mobility model. **(I)** Coefficient of variation of the distribution of total Cdc42T molecules, *CV_patch_* (see Methods), as a function of the number of total GEF molecules. **(J)** Patch centroid diffusivity as a function *CV_patch_* for the Low mobility and High mobility models. Error bars for patch centroid diffusivities are standard errors from the least-squared fit used to compute *D_patch_*. For all other quantities, the error bars are the standard deviation from estimations in 5 independent simulations.

#### B.2. Potential mechanisms for regulating polarity cluster dynamics

During mating, the polarity site transitions from being highly mobile to a more stable state. Therefore, we investigated mechanisms that could be used to regulate patch movement. Our simulation results indicate that increasing GEF abundance can reduce patch mobility (Figs 5E and 5G). We note that increasing *k_6_* or adding Reaction 7 in our model enabled highly mobile clusters at low GEF abundances, but reversing such reaction modifications while keeping low GEF numbers eliminates polarization instead of stabilizing the patch (see for example Fig 5E, *k_6_* = 1 μm^2^/s). After evaluating how all the reactions in the model affect patch mobility (S3 and S4 Figs, Fig 7) we found that the following reactions can substantially and robustly modulate cluster mobility: activation of Cdc42Dm by GEFm (*k_2a_*), activation of Cdc42Dm by Cdc42T-GEF (*k_3_*) and association of Cdc42T with GEFm to form the Cdc42T-GEF complex (*k_4a_*). We note that these reactions all involve the association between two membrane-bound molecules.

**Figure 7.**
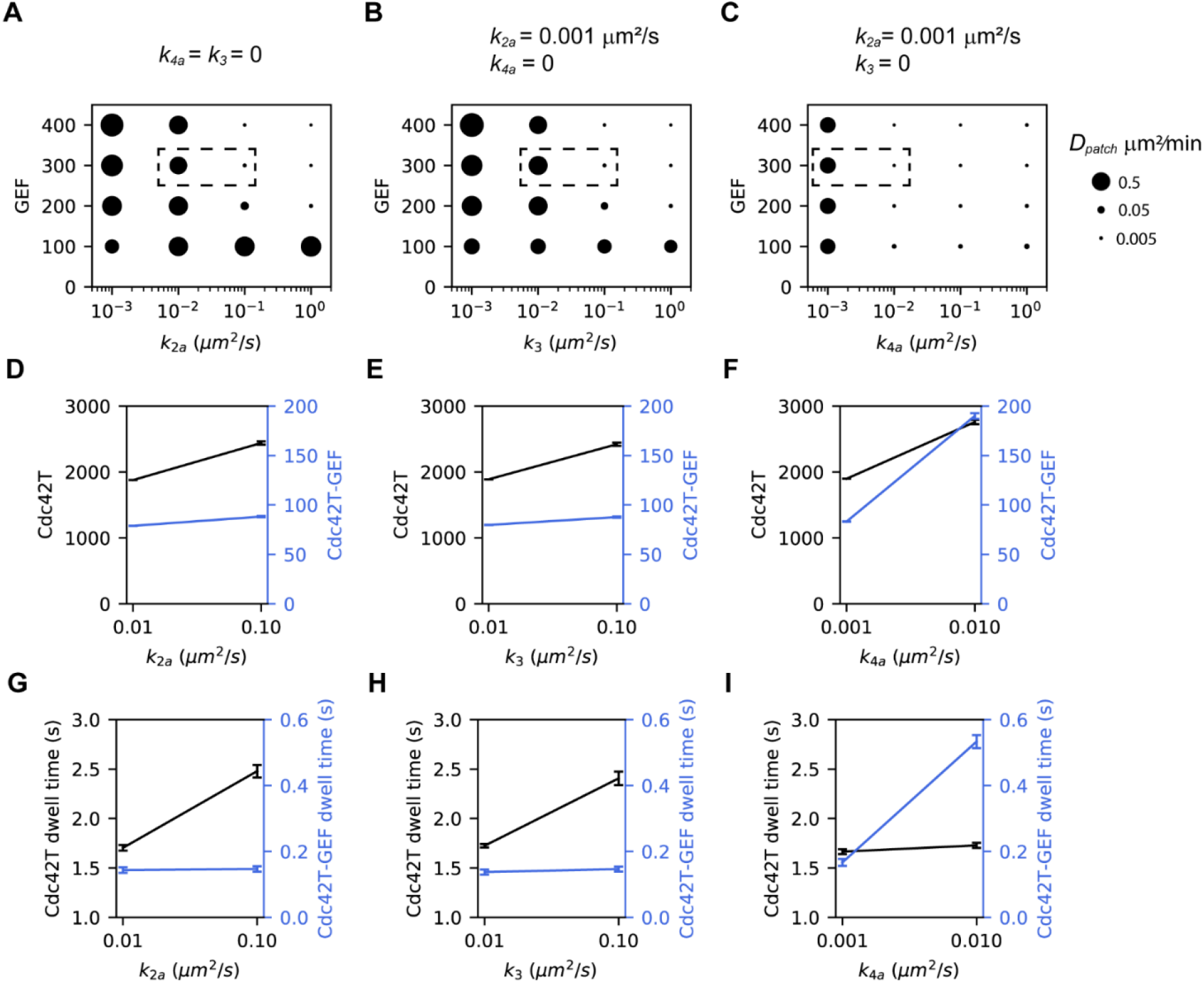
Increasing the association rate constants between molecules at the membrane stabilizes mobile clusters by prolonging the dwell time of polarity factors at the patch. Patch centroid diffusivity (*D_patch_*) as a function of the total number of GEF molecules and *k_2a_* **(A)***, k_3_* **(B)**, *k_4a_* **(C)**. The size of the dots reflects the magnitude of *D_patch_* as indicated in the legend at the bottom-left. For each case, the value of the other two rate constants is shown above the panel. **(D-F)** Abundance of Cdc42T (black) and Cdc42T-GEF (blue) for the corresponding points enclosed by dashed boxes in **(A-C)**. **(G-I)** Dwell times at the membrane of Cdc42T (black) and Cdc42T-GEF (blue) for the corresponding points enclosed by dashed boxes in **(A-C)**. Error bars are the standard deviation from estimations in 10 independent simulations.

To investigate the role of Reactions 2a, 3 and 4a in stabilizing the polarity site, we set the rate constants for the other two either to zero (*k_3_* and *k_4a_*) or a very small value (as for *k_2a_* a minimal value is required to start Cdc42 activation) and varied the rate constant for the remaining reaction and the GEF abundance (Figs 7 A-C). In each case, at a low value of the reaction rate, we observed a mobile cluster even for high GEF abundances. Increasing *k_2a_*, or *k_3_* could stabilize dynamic polarity patches (Fig 7 A,B) for GEF levels of 200 and above, while increasing *k_4a_* stabilized a patch even for GEF = 100. To gain insight into how reaction kinetics modulate patch mobility we computed the total numbers (Figs 7 D-F) and dwell times (Figs 7 G-I) of Cdc42T and Cdc4T-GEF at the patch for the points enclosed in the dashed rectangles in Figs 7 A-C. Each pair of enclosed points correspond to a highly mobile and a stable patch resulting from two different values of the corresponding rate constant. Increasing *k_2a_* or *k_3_* modestly increased the numbers of Cdc42T and Cdc42T-GEF and lengthened the dwell time of Cdc42T. On the other hand, increasing *k_4a_* moderately increased Cdc42T levels but drastically increased the abundance and dwell time of Cdc42T-GEF. These results suggest that increasing the rates *k_2a_* and *k_3_* stabilize the patch mainly by lengthening the dwell time of Cdc42 at the patch, while higher *k_4a_* reduces cluster mobility by increasing both the abundance and dwell time of GEF at the patch.

From Fig 7 A-C it is also apparent that increasing *k_4a_* is a more potent way to stabilize a mobile patch in comparison to increasing *k_2a_* and *k_3_*. This observation led us to investigate in more detail how patch mobility depended on *k_4a_*. With the original parameters of the updated model (including Reaction 7), setting *k_4a_* equal to zero resulted in an increase of more than an order of magnitude in patch mobility for low GEF abundances (Fig 8A). Varying *k_4a_* for GEF=100 resulted in an abrupt change in cluster mobility around a value of 0.003 μm^2^/s (Fig 8B). This transition is likely caused by a synergistic increase in GEF abundance and dwell time at the patch as *k_4a_* increases (Figs 7 C,F,I). After this sharp decrease, the mobility of the patch gradually increased with increasing *k_4a_* and appeared to plateau as the reaction becomes limited by diffusion.

**Figure 8.**
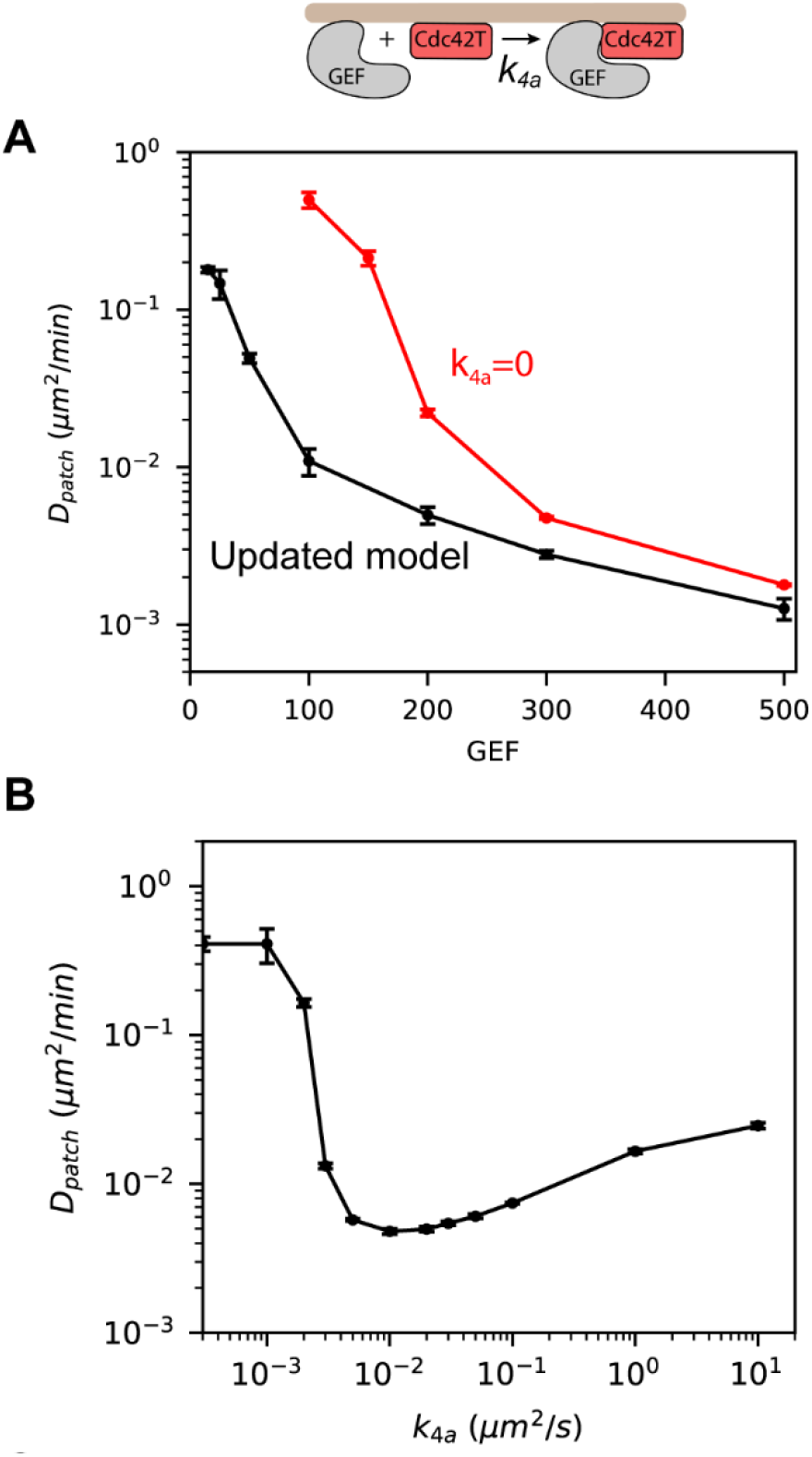
Increasing the rate constant of Cdc42T-GEFm association (Reaction 4a) induces an abrupt change in patch mobility. **(A)** Patch centroid diffusivity as a function of the number of GEF molecules for the model that includes Reaction 7 (Updated model, black) and the same model but setting *k_4a_* = 0 (red). **(B)** Patch diffusivity as a function of *k_4a_* for simulations with GEF = 100. Error bars are standard errors from the least-squared fit used to compute *D_patch_*.

### C. Validation through 3d particle-based simulations

So far, all our results have been obtained using 2D simulations with periodic boundary conditions and assumed the only difference between the membrane and cytosol is the rate at which molecules diffuse. However, real yeast cells are 3D objects with distinct membrane and cytosolic compartments. Therefore, to ensure our results remained valid in 3D, we used Smoldyn [26,27] to perform particle-based simulations on a sphere and we translated 2D parameters to 3D as described in the Methods. Because varying the rate constant *k_4a_* had the largest effect on patch mobility, we chose to validate these results. In agreement with our 2D simulations, low values of *k_4a_* produced a mobile polarity patch and patch mobility decreased rapidly with increasing values of this rate constant (Fig 9A). Indeed, in the 3D particle-based simulations, patch mobility as a function of *k_4a_* showed a similar trend as compared to results from the 2D spatial Gillespie simulations (Fig 9B). For low values of *k_4a_, D_patch_* is high and rapidly decreases as *k_4a_* increases until a minimum value is reached. After that point, patch mobility increases slightly with increasing *k_4a_*. We note, however, that in the 3D particle-based simulations, the transition from a mobile to static patch appears to take place at a slightly higher values of *k_4a_* (between 0.005 and 0.01 μm^2^/s), and for each value of *k_4a_*, *D_patch_* is higher, in comparison with the 2D spatial Gillespie simulations. The good agreement between the full 3D and approximate 2D simulation results validates the use of the more computationally efficient 2D simulations to investigate dynamics of the polarity patch. These results also provide further support for a mechanism for stabilizing the polarity patch by increasing the rate at which membrane-bound GEF and Cdc42T associate.

**Figure 9.**
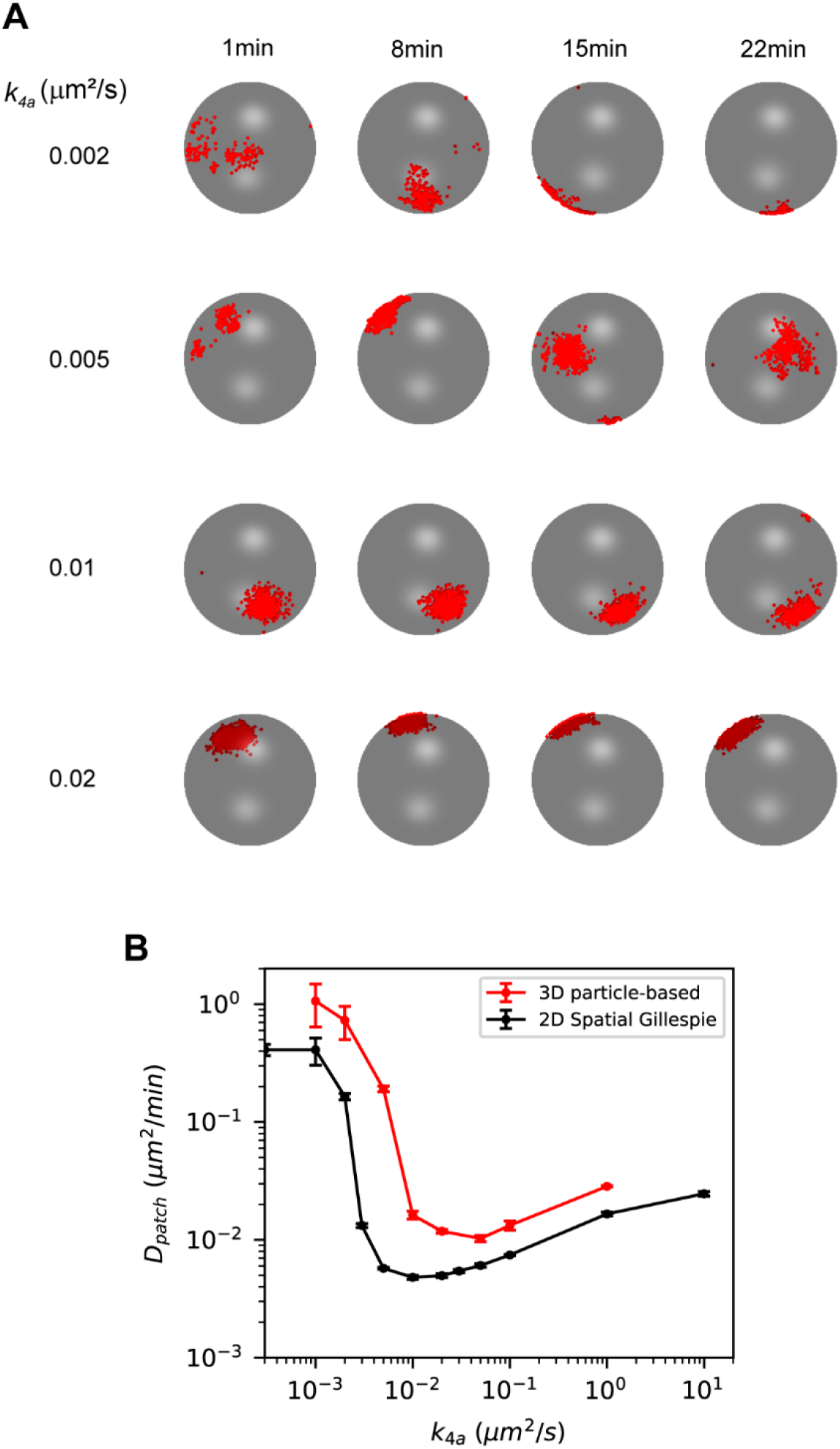
3D particle-based simulations recapitulate 2D spatial Gillespie simulations results from Fig 8. **(A)** Snapshots of 3D particle-based simulations for different values of *k_4a_*. Cdc42T molecules are shown as red dots on a spherical surface representing the cell membrane. Rate constants are estimated from the ones used in Fig 8B as described in the Methods. Parameters are presented in Table 1. **(B)** Patch centroid diffusivity as a function of *k_4a_* for 3D particle-based simulations (red) with GEF = 100. For comparison we also show results from 2D spatial Gillespie simulations in Fig 8. Error bars are standard errors from the least-squared fit used to compute *D_patch_*.

## Discussion

How cells relocate polarity clusters at the cell membrane during different tasks such as migration [30,31], growth [32,33], sporulation [34] and mating [5,35] is a fundamental question that has not been fully understood. By means of computational modeling, we investigated how molecular noise can be exploited to promote lateral mobility of polarity clusters and how cells can regulate cluster mobility. We focused on dynamic polarization observed during the early stages of mating in budding yeast. When haploid cells are presented with pheromone from potential mating partners, the Rho-GTPase Cdc42 forms highly dynamic clusters that move across the membrane and stabilize in the direction of an adjacent cell of the opposite mating type.

### Efficient and accurate stochastic simulations

To investigate the effect of molecular-level fluctuations on cluster mobility, we performed stochastic simulations of the biochemical network of Cdc42 polarization in yeast. Stochastic effects are most accurately captured using particle-based (microscopic) simulations [16,26,27,36–38] or a convergent reaction-diffusion master equation [39]. However, the long-time scales associated with the movement of the polarity site make the use of such simulations computationally prohibitive to perform extensive investigations. We therefore used less accurate, but more computationally efficient simulations based on the spatial Gillespie method. Despite improvements on the accuracy of spatial Gillespie simulations [18–20], we found that simulations lost accuracy in the diffusion-limited regime at high molecular concentrations. To increase the accuracy of such simulations, we implemented concentration-dependent mesoscopic rate constants building on ideas from Yogurtcu et al [21]. An alternative approach that could also provide accurate and computationally efficient results is using hybrid microscopic-mesoscopic simulations [40–45], for example coupling a microscopic formulation of the cell membrane with a mesoscopic representation of the cytosol. Such methods, of course, imply higher implementation complexity.

### Highly dynamic clusters

Relocation of polarity clusters has commonly been explained via negative feedback mechanisms [46]. Negative feedback reactions can destabilize positive-feedback driven clusters resulting in travelling waves [30], or oscillations where the clusters disappear and reappear at different locations of the membrane [35,47]. Directed vesicle delivery is also a form of negative feedback that dilutes the polarity cluster and induces wandering motion [7,13,14]. Here, we document how biochemical noise can induce relocation of polarity clusters without an explicit negative feedback mechanism. Essential for noise-driven cluster motion are low abundances of limiting polarity factors and fast cycling of polarity factors between the cluster and the cytosol. Fast cluster-cytosol cycling can be promoted by reactions where polarity factors are directly recruited to the cluster.

### Regulation of cluster dynamics

During yeast mating, stabilization of highly mobile clusters has been attributed to increased cytosolic levels of the polarity factor Cdc24, the Cdc42 GEF, as pheromone induced MAPK activity triggers its nuclear export [5]. Besides increased GEF abundance, our study suggests that accelerating the kinetics of second-order reactions at the membrane involved in the activation of Cdc42 can stabilize highly dynamic clusters. Increasing the rate constants of such reactions seem to stabilize a mobile cluster by lengthening the abundance and dwell time of polarity factors at the patch. Interestingly, modulating the rate constant of one such reaction, association of membrane-bound Cdc42 and GEF, produced a switch-like transition from a mobile to a stable polarity patch. The Cdc42-GEF interaction, therefore, is a likely target of pheromone induced signaling during regulation of polarity cluster dynamics. Interestingly, this interaction, which in yeast cells is bridged by the scaffold protein Bem1, is thought to be regulated in another context involving cell-cycle control [48].

Further evidence highlights the importance of biochemical events taking place at the cell membrane, and properties of the cell membrane itself in the regulation of polarity cluster dynamics. In fission yeast cells, which also display mobile patches in the early stages of mating, cluster dynamics is known to be under the control of a GAP (Cdc42 inactivator molecule) that localizes at the cell membrane [49]. During spore germination of fission yeast, initial uniform growth is associated with highly dynamic Cdc42 clusters. Upon rupture of the outer spore wall, the clusters stabilize into a single cluster in the direction where rupture takes place, giving rise to directed growth [34]. Other studies have documented that membrane tension [50] and membrane curvature [51,52] can influence cluster stability. Additional mechanisms that may be used by cells to regulate biochemical events at the membrane and control cluster dynamics include crowding of signaling molecules [53,54], restricting the diffusion of molecules with cytoskeletal barriers [11,55,56] and confining molecules into high affinity subdomains [57].

In summary, our results demonstrate the power of using accurate and efficient mesoscopic simulations to inform more detailed, but computationally costly, particle-based simulations. Our studies also provided considerable insight into the mechanisms used by cells to harness random molecular behavior and regulate the dynamic properties of their polarity sites.

## Methods

### Spatial Gillespie simulations

In this coarse-grained approach, space is discretized into grid units, and the state of the system is given by the number of molecules of each species in each grid unit. The system evolves continuously in time according to the reaction-diffusion master equation, which is the spatial extension of the chemical master equation for well-mixed systems. We simulated individual realizations with the Next Subvolume method [17] which is an efficient implementation of the spatial version of the stochastic simulation algorithm [58]. We ran simulations on a square domain of size L discretized with a Cartesian mesh with grid element size *h*. In the spatial Gillespie algorithm, diffusion is treated as a reaction that results in a molecule transitioning from its current location to a neighboring grid unit. If there are *n* molecules of a given species in a particular grid unit, the propensity *k_jump_* for one of those molecules to transition to a neighboring grid unit is *n D/h^2^*, where *D* is the diffusion coefficient. In 2D, there are 4 neighbor cells and the total propensity of jump is 4 *n D/h^2^*. The reaction propensities within a grid unit are estimated in the same way as the well-stirred Gillespie algorithm [58]. For example, the propensity of a second-order reaction for the association of the species A and B, is computed as *k_meso_ n_A_ n_B_* where *k_meso_* is the mesoscopic rate constant and *n*_A_ and *n*_B_ are the numbers of molecules A and B in the particular grid unit. The difference in the versions of the spatial Gillespie methods we use here are the different ways in which *k_meso_* is computed.

### Derivation of the scale-dependent mesoscopic association rate *k_h_* in a 2D system

Hellander et al. [19] derived a mesoscopic rate *k_h_* using the condition that the mean association time for two molecules diffusing in a specified domain in the spatial Gillespie representation is equivalent to the exact result for a microscopic description. The mean association time, τ_micro_, in the microscopic formulation for two molecules with reactive radius ρ diffusing on a disc with radius *R* [18] is:

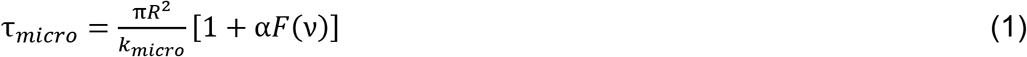

where

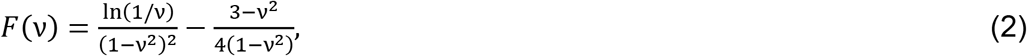

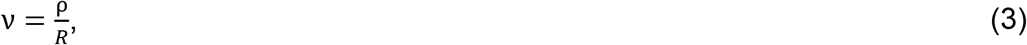

*k_micro_* is the microscopic association rate constant and the parameter α is defined as:

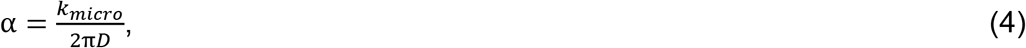

where *D* is the sum of the diffusion coefficients of the two molecules.

The mean association time in the mesoscopic formulation τ_meso_ is estimated for a square domain of length L with square grid units of length *h* [19] as:

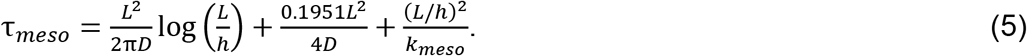

To determine *k_h_*, the condition τ_*meso*_ = τ_*micro*_ is enforced with *L*^2^ = *πR*^2^. This leads to the following expression for *k_h_*:

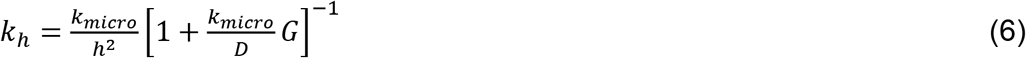

where

Note that *k_h_* can be computed only if:

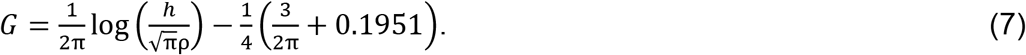

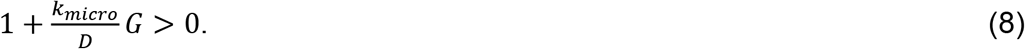

This implies a lower bound on *h*:

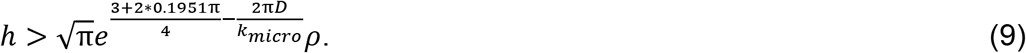

When *h* is below this bound, there is not *k_meso_* for which the equality τ_*meso*_ = τ_*micro*_ holds.

### Mesoscopic dissociation rate *k_h_^d^*

The mesoscopic dissociation rate constant is derived so that the steady state concentrations of a reversible second-order reaction of a two-molecules system are the same in the mesoscopic and microscopic formulations [20]. The steady state is characterized by the ratio of the average unbound time to the total time, which can be computed as the ratio of the mean rebinding time to the sum of the mean rebinding time and the mean dissociation time. Therefore, the condition to be satisfied is:

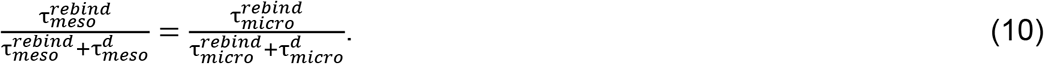

The mean rebinding time in the mesoscopic simulation τ_*meso*_^*rebind*^ was shown to be (*L/h*)^2^/*k_meso_* and a good approximation for the mean rebinding time in the microscopic formulation τ_*micro*_^*rebind*^ in the 2D disk domain is *L*^2^/ *k_micro_*. The mean dissociation times in the mesoscopic (τ_*meso*_^*d*^) and microscopic (τ_*micro*_^*d*^) formulations are respectively 1/*k_meso_^d^* and 1/*k_micro_^d^*. After replacing *k_meso_* and *k_meso_^d^* by *k_h_* and *k_h_^d^* the equilibrium condition then reduces to:

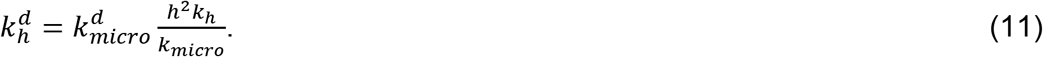

### Mesoscopic concentration-dependent association rate *k*_c_

The mesoscopic concentration-dependent rate *k*_c_ is designed to increase simulation accuracy in systems with high concentrations. In such cases, it is reasonable to ignore the time for two molecules to diffuse into the same grid element, and to focus on the kinetics within a grid element. We define the concentration-dependent rate constant *k*_c_ as:

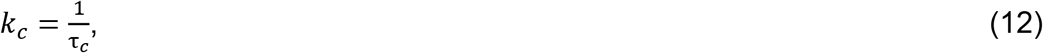

where τ_c_ is the mean association time for two molecules diffusing in an area *A*_c_ corresponding to the mean free area between molecules of the more abundant reactant in the grid element. *A*_c_ is approximated as:

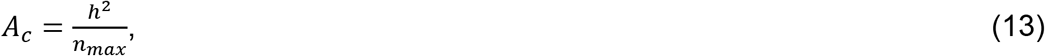

were *n*_max_ is the number of the most abundant species involved in the reaction. τ_c_ is calculated for two molecules diffusing on a disc of area *A*_c_. The concentration-dependent association rate is therefore:

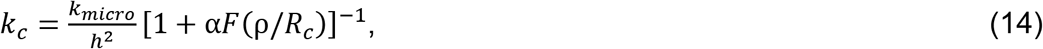

where *F* is the same function defined in equation (2), α = *k_micro_*/(2π*D*), and

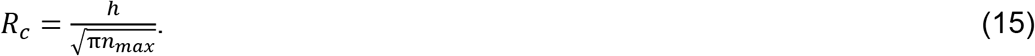

For crowded environments where *R*_c_ ≤ ρ, we set *k_c_* = *k*_micro_. Although technically there is not a lower bound on *h*, for realistic simulations *h* should be greater than ρ.

### Mesoscopic dissociation rate *kc^d^*

We estimate the mesoscopic dissociation rate *k_c_^d^* in a similar way as described above for a two-molecule system under the assumption that the pair of molecules diffuse on a domain of area *A*_c_:

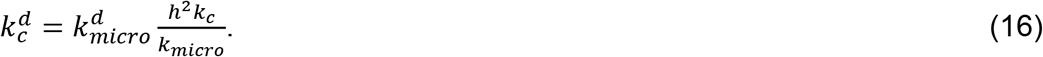

### 2D particle-based simulations

The simulations performed to benchmark our methods (Fig 2) were carried out using our own custom written software. All particle-based polarity simulations (2D and 3D) were performed using Smoldyn [26,27]. 2D simulations were run on a square computational domain with periodic boundary conditions. In particle-based simulations space is continuous and time is discretized in intervals Δ*t*. Molecules are considered point particles and their Brownian motion is simulated with the Euler-Maruyama method: If *x*(*t*), *y*(*t*) are the position coordinates of a given particle at time *t* moving in a 2D domain, the position coordinates at time *t* + Δ*t* are calculated as:

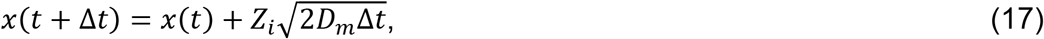

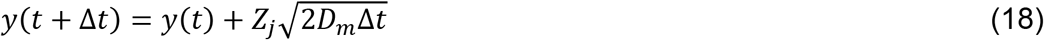

where *Z_i_*, and *Z_j_* are independent random numbers drawn from a standard normal distribution, and *D_m_* is the diffusion coefficient. Every time step, the new positions of all the particles are calculated.

Bimolecular reactions occur with probability P = 1-exp(-*λ* Δ*t*) during a time interval Δ*t*, which can be approximated as P = *λ* Δ*t* for small Δ*t*, when two reactants are within a distance ρ (reactive radius).

The reaction probability for a first-order reaction with rate constant *k_i_* during an interval Δ*t* is P_*i*_ = 1-exp(-*k_i_* Δ*t*), which can be approximated as P_*i*_ = *k_i_* Δ*t* for small Δ*t*. When a dissociation event for two molecules in a complex occurs, one molecule is set at the position previously occupied by the complex and the second is placed at distance ρ apart from the first, with the orientation chosen randomly from a uniform distribution.

Simulations of simple reversible and irreversible reactions (Fig 2) were performed with Δ*t* ≈ (0.1ρ)^2^/(4*D*_tot_) with *D*_tot_=2*D_m_*. The parameters of such simulations are given in the captions of the corresponding figures.

For simulations of the polarity establishment model we used the software Smoldyn [26,27]. The parameters for the polarity establishment model are given in Table 1. We present the reaction parameters as microscopic rate constants *k_micro_*. For second-order reactions, the input to Smoldyn is the reaction probability P = *λ* Δ*t*, and *λ* is related to *k_micro_* as:

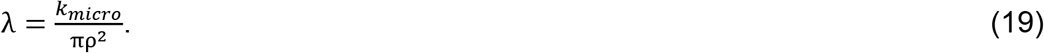

This equality follows as *k_micro_* quantifies the reaction probability rate per unit area (*λ*/πρ^□^) for a pair of molecules *A*, *B* that are within the reactive distance ρ:

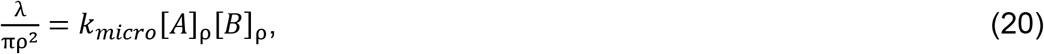

### Reactions of the polarization model

|  |  |
| --- | --- |
| $GEF_c \rightarrow GEF_m$ | R-1a |
| $GEF_m \rightarrow GEF_c$ | R-1b |
| $GEF_m + Cdc42D_m \rightarrow GEF_m + Cdc42T$ | R-2a |
| $Cdc42T \rightarrow Cdc42D_m$ | R-2b |
| $Cdc42T-GEF + Cdc42D_m \rightarrow Cdc42T-GEF + Cdc42T$ | R-3 |
| $Cdc42T + GEF_m \rightarrow Cdc42T-GEF$ | R-4a |
| $Cdc42T-GEF \rightarrow Cdc42T + GEF_m$ | R-4b |
| $Cdc42D_c \rightarrow Cdc42D_m$ | R-5a |
| $Cdc42D_m \rightarrow Cdc42D_c$ | R-5b |
| $Cdc42T + GEF_c \rightarrow Cdc42T-GEF$ | R-6 |
| $Cdc42T-GEF + Cdc42D_c \rightarrow Cdc42T-GEF + Cdc42T$ | R-7 |

In the above reactions, GEFc and GEFm are cytosolic and membrane bound GEF, respectively. Cdc42Dc and Cdc42Dm are cytosolic and membrane-bound inactive Cdc42 (Cdc42-GDP), respectively. Cdc42T is membrane-bound active Cdc42 (Cdc42-GTP). Cdc42T-GEF is the membrane-bound complex of Cdc42T and GEF.

### Rate constants for the 2D stochastic polarization model

The rate constants for the 2D model in Fig 3 A, presented in Table 1, were adapted from [13] which is a modified version of the model in [23]. That model is based on reaction-diffusion equations and is therefore a macroscopic representation. On the other hand, our stochastic simulations are parameterized with microscopic rate constants. For first-order and second-order reaction-limited reactions, the macroscopic rate constants from the model in [13] can be used directly in our simulations. However, for 2D diffusion-influenced reactions the conversion is more complicated [16]. For the purposes of this work, we used the macroscopic rate constants in [13] as a first approximation for the microscopic parameters and performed careful studies varying the rate constants over several orders of magnitude.

While the cell cytosol is a 3D compartment, in our 2D simulations the cytosol and the membrane are juxtaposed two-dimensional domains. This is a computationally efficient representation that neglects cytosolic gradients perpendicular to the membrane since, diffusion at the cytosol is fast compared to the timescale of reactions. To obtain equivalent rate constants for this purely 2D system we scaled cytosolic concentrations in the rate equations in [13] as:

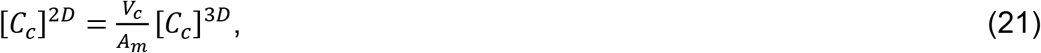

where [C_c_]^3D^ is the molar concentration of the cytosolic component C_c_ in the original equations, [C_c_]^2D^ is the concentration of C_c_ in the 2D cytosol, V_c_ is the volume of the cell cytosol and Am is the membrane area.

In [13], concentrations at the membrane are expressed in molar units assuming that the membrane was a volumetric compartment with thickness *Th*. We therefore also scaled concentrations of species at the membrane as:

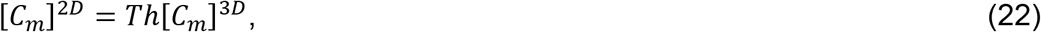

where [C_m_]^2D^ is the concentration of the membrane-bound species C_m_ in units of mass/area, and [C_m_]^3D^ is the molar concentration of C_m_. From the scaled reaction-diffusion equations we obtain the scaled 2D reaction rate constants *k^2D^* in terms of the rate constants *k^3D^* in [13]:

- Rate constants for first-order reactions that involve a transition from the cytosol to the membrane (R-1a and R-5a) are scaled as:

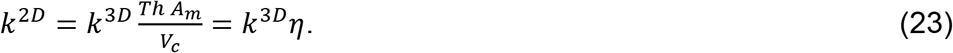
- Second-order rate constants for reactions taking place at the membrane (R-2a, R-3, R-4a) are scaled as:

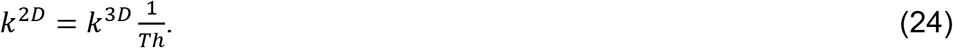
- Second-order rate constants for reactions in which a cytosolic species reacts with a membrane-bound species (R-6, R-7) are scaled as:

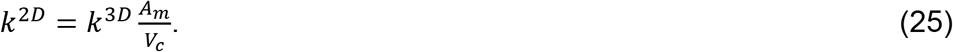
- The rate constants for first-order reactions in which the reactant and the product are bound to the membrane (R-2b and R-4b) are unchanged.
- Reactions 1b and 5b are the reverse of reactions 1a and 5a respectively, and to obtain the 2D rate constants it is necessary to multiply the corresponding volumetric rate constants by a factor of 1/η. However, that factor cancels out with a factor of η present in the reaction-diffusion equations used in [13,23], which takes into account the difference between the cytosol volume and the effective volume of the membrane. Therefore, *k_1b_* and *k_5b_* are unchanged.

The resulting 2D rate constants are presented in Table 1 using molecules as the unit of mass and μm as the unit of length.

We note one significant difference between the model parameters used here, and those used by McClure et al. We used a smaller number of total Cdc42 molecules based on the work of Watson et al. [59]. For our particle-based simulations, we used a smaller reaction radius ρ than in our previous publication [16] to make this value closer to the size of the reacting proteins. This modification reduced the rate of Cdc42 activation, affecting polarization. To compensate for this effect, we increased the rate constants of Reactions 5a, 5b and 6 by a factor of 10.

### 3D particle-based simulations

3D simulations were performed using Smoldyn [26,27]. Membrane-bound species diffuse on the surface of a sphere with radius R and cytosolic components diffuse within the interior of the sphere. The parameter values used in the particle-based 3D simulations are given in Table 1.

The reaction rate constants were obtained from the 2D model. Rate constants for reactions that take place exclusively in the membrane do not need to be modified. These are Reactions 1b, 2a, 2b, 3, 4a, 4b and 5b.

The rate constants for reactions in which a cytosolic species binds the membrane or a membrane-bound molecule (Reactions 1a, 5a, 6, and 7) are estimated from the 2D model using the scaling introduced in the Methods subsection “Rate constants for the 2D stochastic polarization model” ignoring the factor *Th*. With that scaling these 3D rate constants are obtained by multiplying the corresponding 2D rate constants by the factor *V_c_*/*A_m_*.

Although we report the microscopic rate constant *k_micro_* for all reactions, in Smoldyn, second-order reactions are parameterized with the reaction probability P = *λ* Δ*t*. For reactions where both reactants are at the membrane, the relation between λ and *k_micro_* is the same as in the 2D simulations.

For second-order reactions where a cytosolic molecule reacts with a membrane-bound molecule and the product is at the membrane (Reactions 6, 7) λ is related to *k_micro_* as:

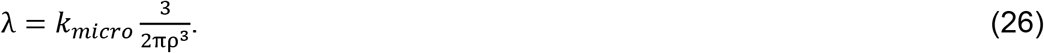

This is the case as *k_micro_* determines the reaction probability rate per unit area (λ/πρ^2^) for a membrane-bound molecule *A^m^* and a cytosolic molecule *B^c^* that are within a distance ρ:

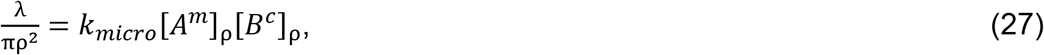

where [*A^m^*]_ρ_ = 1/(πρ^2^), and [*β^c^*]_ρ_ = 3/(2πρ^3^) which is the concentration of a molecule in a half sphere of radius ρ.

### Quantification of clustering with H(*r*)

Molecule clustering is commonly assessed by computing the cumulative distribution function (cdf) of inter-particle distances *r*, which properly normalized is known as Ripley’s K function K(*r*). This function is compared between the system of interest and a reference situation where particles are uniformly distributed (null hypothesis). In particular, H(*r*) is constructed by subtracting from K(*r*) the corresponding function for a uniform distribution of particles. Therefore, H(*r*) = 0 if the frequency of inter-particle distances within a separation *r* is the same as that of uniformly distributed molecules, indicating that there is no clustering. H(*r*) < 0 indicates dispersion; and H(*r*) > 0 indicates clustering or aggregation. The value of *r* where H(*r*) is maximum provides an estimate of the cluster length-scale [24]. We computed H(*r*) as described in [25].

### Mean squared displacement (MSD) and effective polarity cluster diffusivity (*D_patch_*)

After a polarity cluster has formed, the distribution of active Cdc42 is translated to the center of the domain to reduce border effects, and the centroid of the distribution is recorded every 1min. The centroid of the patch is calculated accounting for the toroidal geometry of the domain resulting from the periodic boundary conditions. The mean squared displacement (MSD) for a particular time interval Δ*t_i_* is computed from all centroid trajectories over time intervals of length Δ*t_i_* from multiple simulations. We discarded centroid jumps over the domain boundary by breaking trajectories containing jumps larger than a maximum jump *max_jump_* into sub-trajectories containing only jumps smaller than *max_jump_*. We empirically set *max_jump_* = 6μm from visual inspection of centroid trajectories. Each MSD curve is obtained from 50 simulations of 1hr each except for S4 Fig where we used 5 simulations. The effective diffusivity of the polarity cluster (*D_patch_*) can be obtained by fitting the equation MSD(Δ*t_i_*) = 4*D_patch_* Δ*t*_*i*_^β^ to the MSD data, where Δ*t_i_* is a particular time interval, and a reflects the degree of anomalous diffusion. In practice we took logarithms to the data and fit the equation log(MSD(Δ*t_i_*)) = *log(4D_patch_*) + β log(Δ*t_i_*) using only the first data points that showed a linear behavior in a log-log plot. The MSD during the smallest interval computed (1min) reflects rapid variations in the position of the centroid within the polarity cluster and do not contribute to the long scale displacement of the distribution (see for example Fig 5B). We therefore subtracted MSD(1min) to all MSD data before estimating *D_patch_*. MSDs in the 3D particle-based simulations were computed from geodesic displacements of the cluster on the spherical surface.

### Coefficient of variation of the distribution of active Cdc42T: *CV_patch_*

*CV_patch_* was computed as a weighted average over space of the local coefficient of variation over time of the amount of Cdc42T. The average over space is weighted by the mean local abundance of Cdc42T:

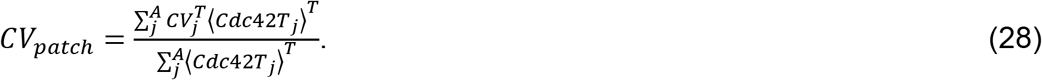

Here *CV_j_^T^* is the mean divided by the standard deviation over a time interval T of the amount of Cdc42T at location *j*. <*Cdc42T_j_*> is the average amount of Cdc42T at location *j* over a period of time *T*. The space average is computed over the whole simulation domain (A) as Cdc42T is mainly located at the polarity site. The time averages are computed over a short period *T* = 1 min to ensure the mean distribution of Cdc42T does not relocate significantly. One estimation of *CV_patch_* is obtained from a single simulation that has reached steady state with samples taken every second to compute time averages. In Figs 6 G,H we plotted the mean and standard deviation (error bars) from 5 independent measurements of *CV_patch_*.

### Dwell times at the patch

To compute the dwell time of Cdc42 at the patch, we introduced in the model additional tagged Cdc42 species (Cdc42_m_^tagged^, Cdc42T^tagged^, Cdc42T^tagged^-GEF) that have the same behavior as the untagged versions, except that if Cdc42_m_^tagged^ jumps from the membrane to the cytosol it converts into untagged Cdc42Dc. Simulations are initialized with no tagged species, and are run until the distribution of polarity factors reaches steady state. At this point, Cdc42m, Cdc42T, Cdc42T-GEF are converted into the tagged versions in a region surrounding the polarity patch and we record the decay in the amount of the tagged molecules. The same idea is used to estimate the dwell time at the patch of GEF (introducing GEFm^tagged^ and Cdc42T-GEF^tagged^). For Cdc42, we ignored the initial rapid decay coming from membrane detachment of inactive Cdc42. The dwell time at the patch is obtained by fitting an exponential decay function to the data. We reported the mean and standard deviation (error bars) from 10 or 30 independent measurements of the dwell time.

## Code availability

The code used to generate all the data and figures is available at: https://github.com/samuramirez/stochastic-exploratory-polarization

## Acknowledgements

We thank members of the Elston lab for helpful discussion throughout the project. This work was supported by NIH/NIGMS grants R35 GM127145 to T.C.E. and R35 GM122488 to D.J.L.

## Supplemental material

### Molecular abundance fluctuations from spatial Gillespie simulations of irreversible and reversible association reactions

**Figure S1.**
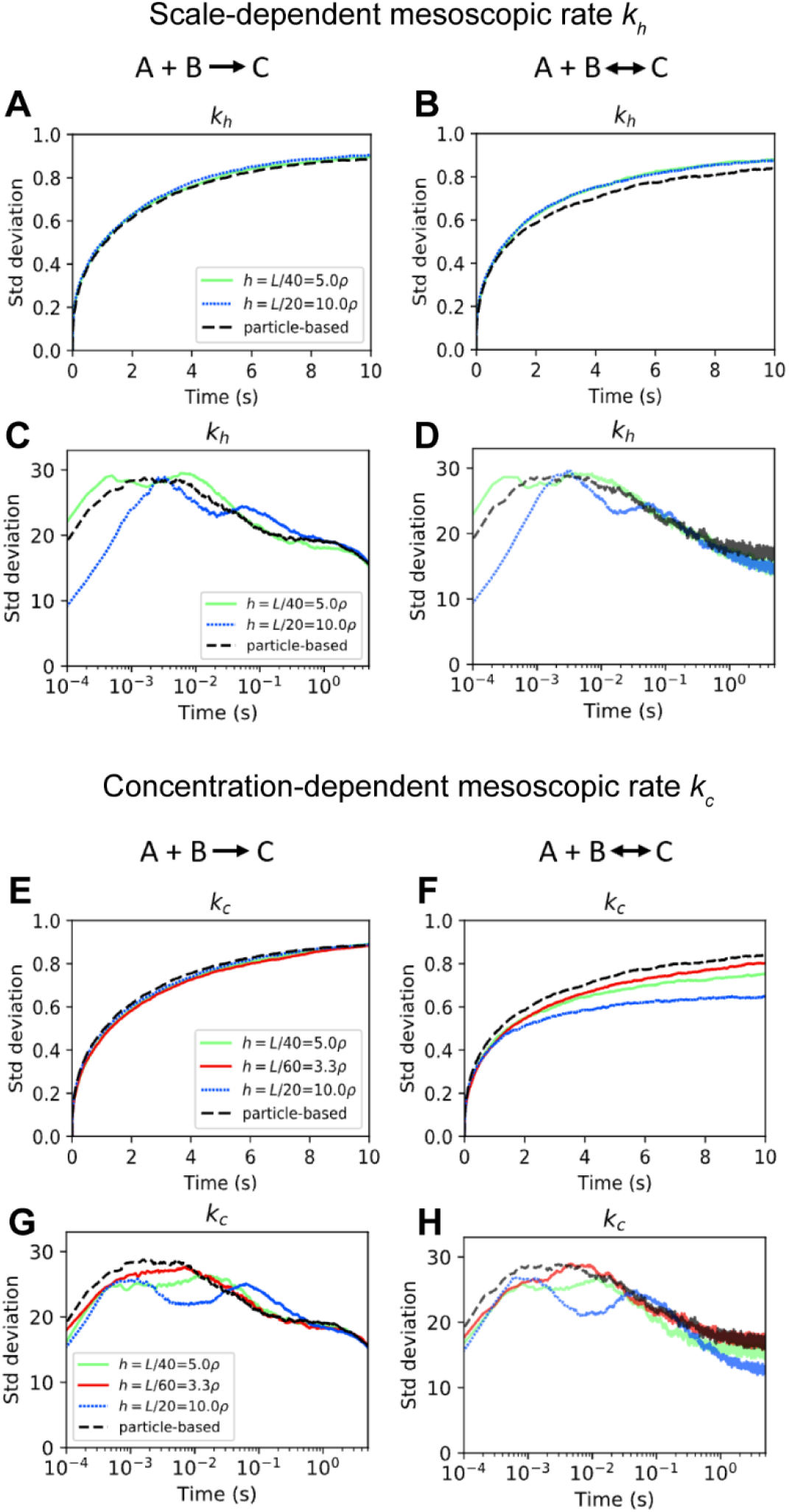
Fluctuations around the mean for spatial Gillespie simulations of the reactions A+B → C and A+B ↔ C using the mesoscopic rates *k_h_* and *k_c_* are compared particle-based results. The mean as a function of time is shown in Fig 2 of the main text. We present the standard deviation of total number of species A as a function of time. **(A-D)** show spatial Gillespie simulations using the mesoscopic rate *k_h_* with initial low abundance of reactants in **(A,B)** (total A = total B = 5, total C = 0 at t = 0) and initial high abundance (total A = total B = 5000, total C = 0 at t=0) in **(C-D)**. **(E-H)** show corresponding simulations to **(A-D)** but using the mesoscopic rate *k_c_*. In all the simulations, the degree of diffusion control is *k_micro_/D_tot_* = 50, with *D_tot_* = 2*D* and *D* = 0.0025μm^2^/s. The size of the domain is *L* = 1μm, ρ = 0.005μm. For the reversible reaction A+B ↔ C, the microscopic dissociation rate constant *k^d^_micro_* is 1/s in panels **(B, F)**, and 10/s in panels **(D, H)**.

### Parameter exploration to enable polarization with low GEF numbers

We varied different rate constants with goal of finding parameter sets that enable polarization with low GEF numbers. To evaluate if the distribution of active Cdc42 was polarized for a given parameter set we quantified clustering with H(*r* = 1.1 μm) (see Methods and Main Text). In most cases, a value of H(*r* = 1.1 μm) > 1.5 indicated there was polarization, but when the numbers of active Cdc42 are low the value of H can fluctuate substantially. Therefore, we also used the total amount of active Cdc42 as an additional criteria for polarization. For cases of highly variable clustering, amounts of active Cdc42 close to zero indicated no polarization.

Parameter changes that resulted in polarization with 100 GEF molecules or less were decreasing *k_2b_*, decreasing *k_4b_*, increasing *k_5a_* and increasing *k_7_*.

**Figure S2.**
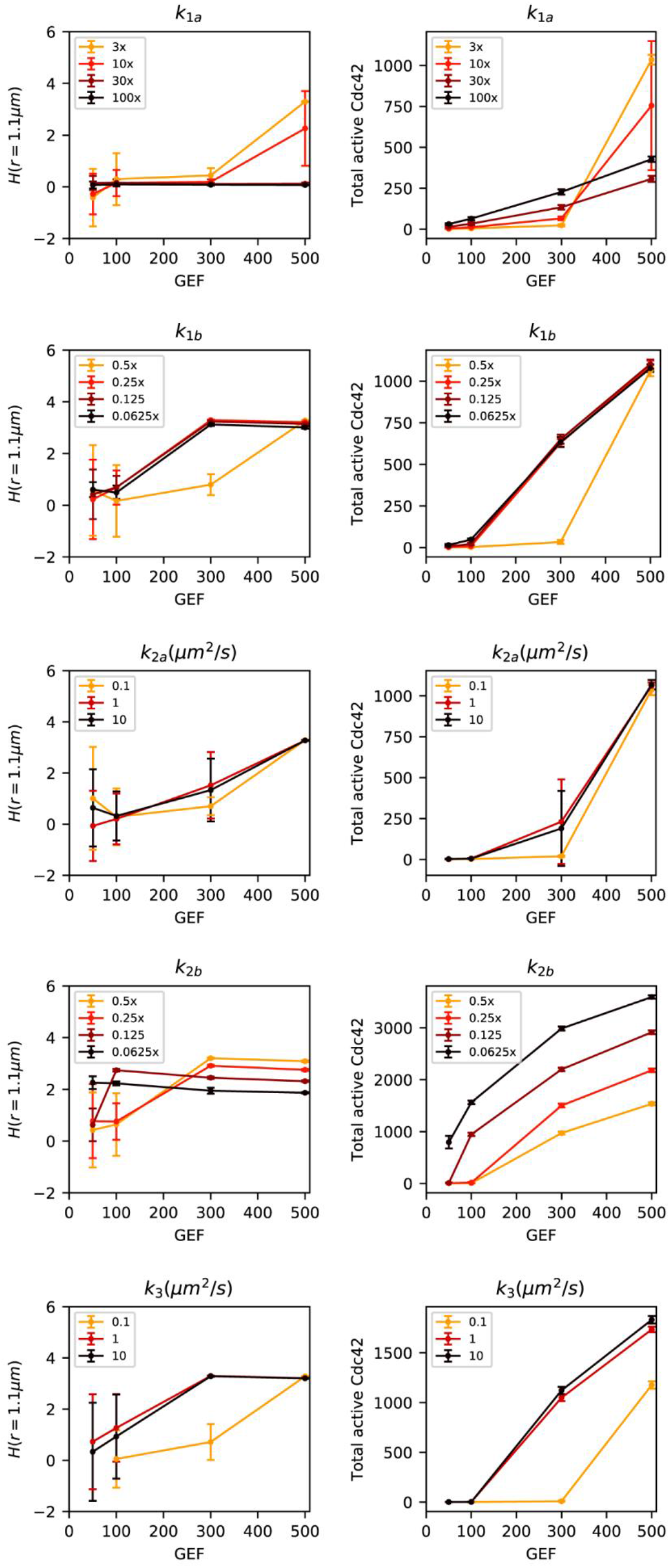
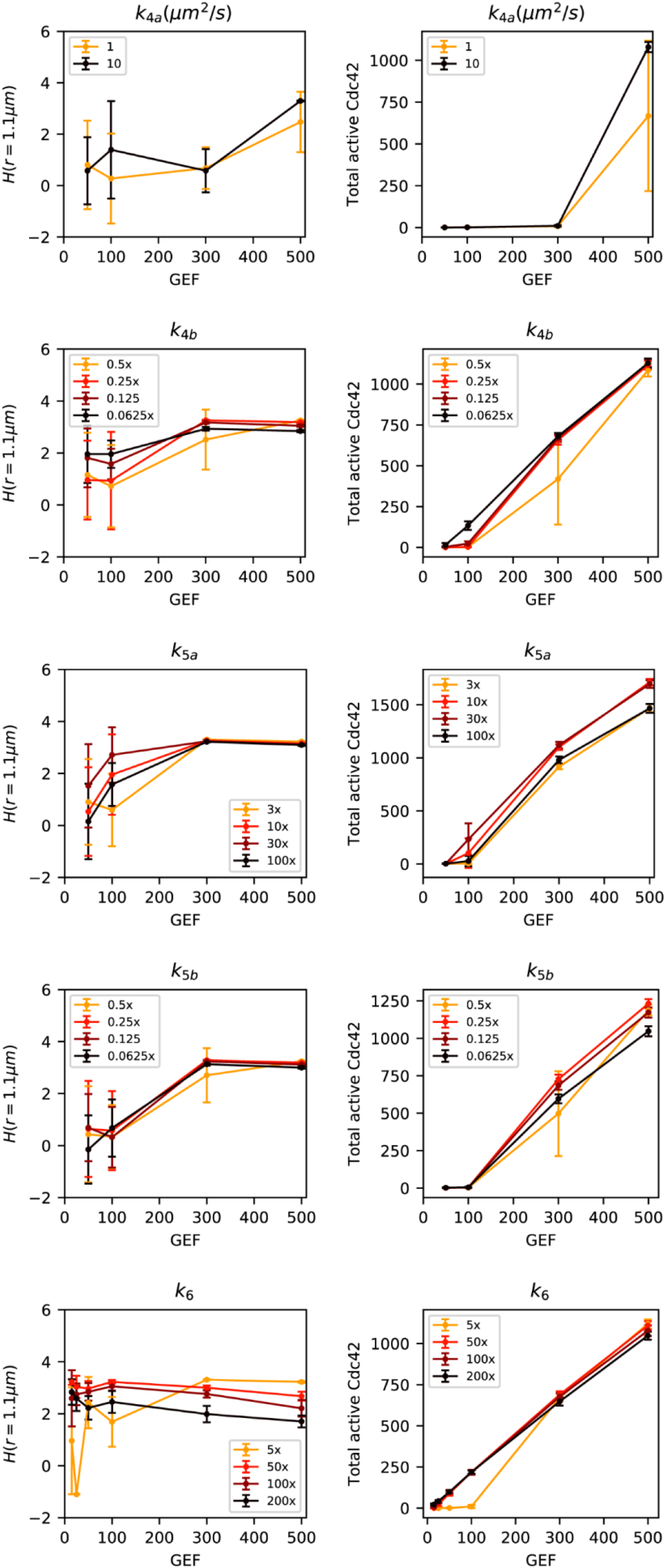
Plots of H(*r* = 1.1 μm) (left panels) and total active Cdc42 (right panels) as a function of GEF molecules for different values of various model parameters. In each panel the rate constant in the title of the figure is varied as indicated in the legend. For each parameter set, the simulations were initialized with an unpolarized random distribution with all GEF and Cdc42 in the cytosol and 10 min were simulated to provide enough time for the system to polarize. Mean and standard deviation (error bars) were calculated sampling every 30s for the last 5min of each simulation with data from 3 independent simulations.

### Testing how different rate constants affect patch mobility

As described in the main text the parameter *k_4a_* has a strong influence on patch mobility. In our initial parameterization, the value of *k_4a_* is relatively high, making the patch stable and obscuring the effect of other parameters. For example, changing *k_3_* from 1 μm^2^/s to zero (in the updated model that includes Reaction 7) does not affect patch mobility significantly (S3 A Fig). However, when the value of *k_4a_* is set to zero, for a range of GEF abundances, changing *k_3_* from its original value of 0.07 μm^2^/s to zero increases patch mobility significantly (S3 B Fig). In S3 C Fig, for 300 GEF, we show that the change in patch mobility takes place gradually for different values of *k_3_*.

To test the effect of patch mobility of other rate constants, we started with the following parameterizations of the updated model (including Reaction 7) which show different levels of patch mobility:

- *k_4a_* = 2 μm^2^/s with GEF numbers of 50 (high mobility) 100 (medium mobility) and 300 (low mobility). These results are shown on the left column of S4 Fig.
- *k_4a_* = 0 with GEF numbers of 150 (high mobility) 250 (medium mobility) and 400 (low mobility). These results are shown on the right column of S4 Fig.

In S4 Fig each row corresponds to a particular rate constant tested as indicated on the x axis. We show data for all rate constants except for *k_2a_* and *k_4a_* which are presented in the Main Text and *k_3_* which is evaluated in S3 Fig.

Varying the value of most of the parameters did not substantially affect patch mobility. Modifying *k_5b_* (the rate constant for Cdc42D to transition from the membrane to the cytosol) showed a moderate effect on patch mobility for *k_4a_* = 2 μm^2^/s. The effect was less robust when *k_4a_* = 0, in this case changing *k_5b_* tended to destroy polarity (missing points in the curves).

**Figure S3.**
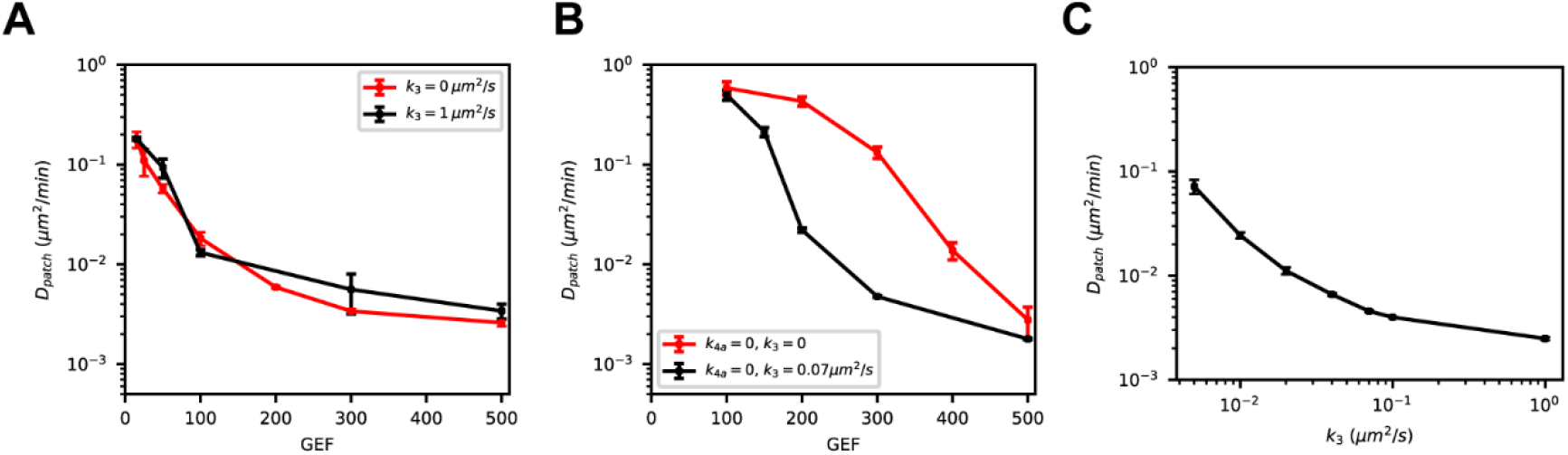
The rate constant *k_3_* affects patch mobility when *k_4a_* = 0. (**A**) Effective diffusivity of the patch (*D_patch_*) as a function of total available GEF in the updated model (including Reaction 7) for *k_3_* = 0 and *k_3_* = 1 μm^2^/s. (**B**) Similar to (**A**) except with *k_3_* = 0 and *k_3_* = 1 μm^2^/s keeping *k_4a_* = 0. (**C**) Effective diffusivity of the patch (*D_patch_*) as *k_3_* is varied with 300 GEF molecules. Error bars are standard errors from the least-squared fit used to compute *D_patch_*.

**Figure S4.**
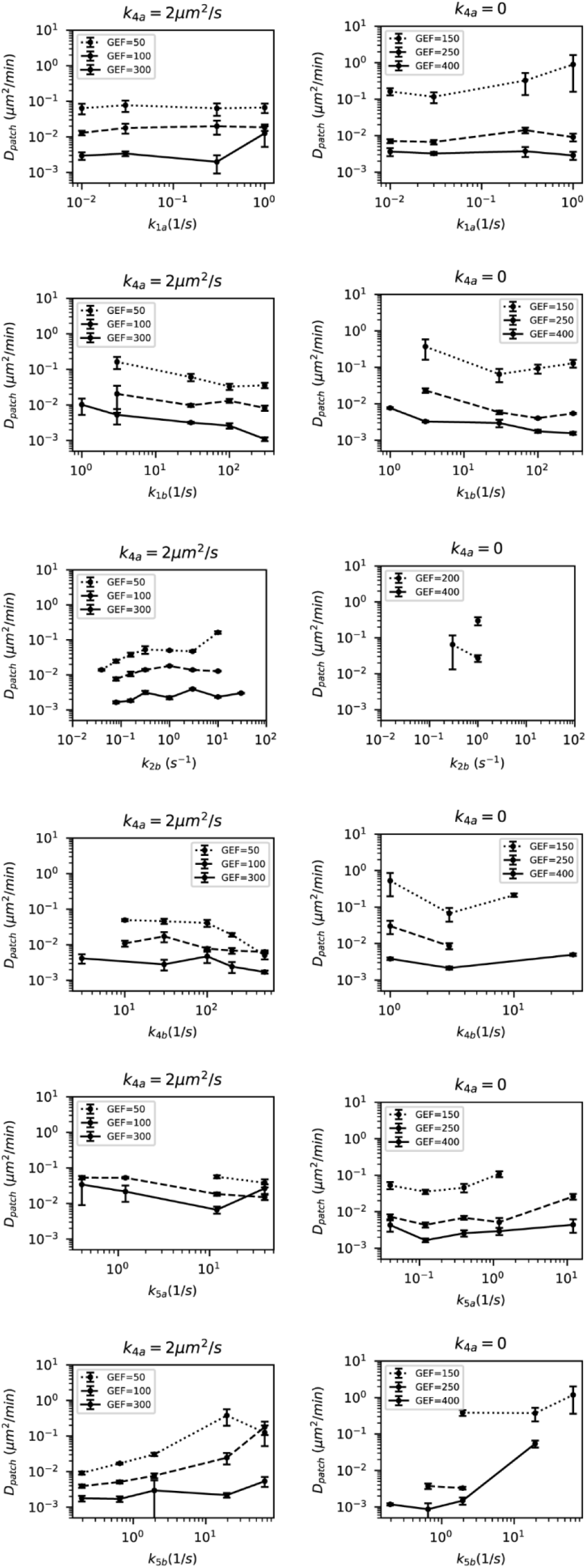
Effective diffusivity of the polarity patch as a function of different parameters (indicated in the x-axis). The panels on the left are for *k_4a_* = 2 μm^2^/s and for the ones on the right *k_4a_* = 0. Simulations were run with different GEF abundances as indicated. Missing points in each panel correspond to simulations that did not show robust polarization. Each data point was obtained from 5 simulations of 3600s each as described in the Methods. Error bars are standard errors from the least-squared fit used to compute *D_patch_*.

